# Reversible solidification of fission yeast cytoplasm after prolonged nutrient starvation

**DOI:** 10.1101/368076

**Authors:** Maria B. Heimlicher, Mirjam Bächler, Chieze Ibeneche-Nnewihe, Ernst-Ludwig Florin, Andreas Hoenger, Damian Brunner

## Abstract

Cells depend on a highly ordered organization of their content and they must develop strategies to maintain the anisotropic distribution of organelles during periods of nutrient shortage. One of these strategies, observed in bacteria and in yeast cells with acutely interrupted energy production, is to solidify the cytoplasm. Here, we describe a different type of cytoplasm solidification that occurs in fission yeast cells having slowly run out of nutrients after multiple days of culturing. It provides the most profound reversible cytoplasmic solidification of yeast cells described to date. Our data suggest the involvement of a matrix with a certain mesh size that immobilizes cellular components in a size-dependent manner. We provide experimental evidence that cells need time, intrinsic nutrients and intrinsic energy sources to enter this state in the absence of external sources. Such cytoplasmic solidification may provide a robust means to protect cellular architecture in dormant cells.

## Introduction

Cell function and survival require a highly ordered, cell type-specific organization of the cytoplasmic content. How cells generate and maintain a cell type-specific spatial organization is central to understanding the living state. It involves the asymmetric distribution of material, mediated by active transport of key regulatory components, which counteracts the entropic activity of diffusion. This occurs in a highly crowded and dynamic environment, with spatially dispersed constituents ranging in size from small ions and metabolites to macromolecular complexes and to large, complex structures like the cytoskeletal networks and organelles. Their sub-diffusive motion is influenced by macromolecular crowding, specific and unspecific interactions, and the polymer networks of the cytoskeleton (Luby-Phelps et al. 1986; Tolić-Nørrelykke et al. 2004; Weiss et al. 2004; Wirtz 2009). Consequently, the cytoplasm was described as a complex viscoelastic fluid, a gel-like material, or a colloidal liquid at the transition to a glass-like state (Luby-Phelps et al. 1986; Tolić-Nørrelykke et al. 2004; Mitchison et al. 2008; Fels et al. 2009; Miermont et al. 2013; Moeendarbary et al. 2013; Parry et al. 2014; Grygorczyk et al. 2015). The maintenance of such a complex, anisotropic cellular architecture and its remodeling in response to environmental changes requires a constant input of energy. Organisms have adopted a variety of strategies to cope with situations in which energy is limited. For example, when nutrients become limiting for growing and dividing cells they can exit the cell division cycle and enter quiescence, similar to when they start differentiating (Coller et al. 2006). Such nutrient starvation-induced quiescence reverses as soon as nutrients become available again (Yanagida 2009; Lennon & Jones 2011). In extreme cases of energy shortage, cells can enter a dormant state with little or no energy consumption (Lennon & Jones 2011). This mainly involves the formation of specialized cell types such as spores and seeds, some of which can survive for centuries. Some animal species, such as tardigrades or hibernating mammals, manage to adopt dormant states by massively down regulating cellular metabolism (Guppy & Withers 1999; K. B. Storey & J. M. Storey 2007; Reuner et al. 2009). In most cases, it is not known to what extent, if at all, the low energy consumption during such dormant states allows cells to maintain their favored cytoplasmic organization. One way to preserve cellular architecture during an extended dormancy period is to reduce water content. This increases macromolecular crowding, which can trigger a transition into a glass-like state of the cytoplasm constraining the motion of its cytoplasmic components. This mechanism was shown to operate in bacterial spores, in plant seeds, in metabolically inactive bacteria, and in budding yeast cells acutely depleted of energy (Sun & Leopold 1997; Cowan et al. 2003; Dijksterhuis et al. 2007; Parry et al. 2014; Joyner et al. 2016; Munder et al. 2016). Alternatively, cells replace water with high amounts of carbohydrates, possibly providing cytoplasmic vitrification to prevent harmful fusion events (Soto et al. 1999; Elbein et al. 2003). Furthermore, quiescent cells were shown to introduce structural changes to cytoplasmic components following energy depletion, with many metabolic enzymes and other proteins forming transient assemblies (Sagot et al. 2006; Laporte et al. 2008; Narayanaswamy et al. 2009; Noree et al. 2010; O’Connell et al. 2012; Petrovska et al. 2014). These assemblies were suggested to inactivate enzymatic function and to serve as storage depots. As a collective, they were proposed to form higher-order structures that mediate a solid-like state of the cytoplasm (Munder et al. 2016).

When nutrients become growth limiting for cells of the fission yeast *Schizosaccharomyces pombe*, they will mate and produce dormant spores (Tanaka & Hirata 1982; Egel 1989). Cells lacking a mating partner will enter a quiescent state. So far, research on fission yeast starvation has largely focused on the physiology of nitrogen-starved cells or cells acutely depleted of glucose (Costello et al. 1986; Yanagida et al. 2011; Saitoh & Yanagida 2014; Oda et al. 2015; Joyner et al. 2016). We describe fission yeast cells that have slowly run out of nutrients after several days of culturing. These cells show a drastic solidification of their cytoplasm, which we termed “cytoplasmic freezing” (CF). In this state, the diffusive motion of lipid droplets and other structures observable by phase contrast or DIC microscopy is almost completely restricted. Cytoplasmic solidification in CF cells is more profound and robust in contrast to other known liquid- to solid-like transitions (Joyner et al. 2016; Munder et al. 2016). We find no evidence for CF being caused by increased macromolecular crowding. Instead, our data suggest that CF may be caused by the formation of some sort of global cellular matrix with a certain mesh size that stably traps the bigger cytoplasmic components whereas small molecules still are able to diffuse freely.

## Results

### Starved cells have 2 intracellular immobilization states

After 2 days of culture under standard conditions, cells of the fission yeast *S.pombe* suffer glucose starvation (GS), arrest growth and enter a quiescent state (Makushok et al. 2016). Imaging cells during the following days of GS, revealed a striking decrease in mobility of virtually all subcellular structures visible by light microscopy, which ended with their seemingly complete immobilization (Movie 1; Methods). An exception were small particles that erratically moved within vacuoles throughout starvation. We quantified this change in mobility over time by tracking the motion of lipid droplets (LDs), which are clearly distinguishable when imaging cells with DIC, at each culturing day (Methods). In our experiments, culturing day 2 will be referred to as starvation day 2 (SD2), culturing day 3 as starvation day 3 (SD3) etc. In exponentially growing cells the lipid droplets are dispersed throughout the cell and dynamically move throughout the entire intracellular space (Figure 1A, Movie 1). On SD2, LDs frequently accumulated into 1–2 grape-like structures displaying visible motion, although their overall position within cells remained fairly constant (Figure 1 – Figure supplement 1A). On SD3-SD4 LDs showed similar dynamicity but an increasing number of cells had fewer, larger LDs, presumably a product of fusion within the grape-like structures (Figure 1 – Figure supplement 1A). By SD5 and SD6 this LD phenotype was prominent in a large fraction of cells. Strikingly, a complete absence of motion was observed for all LDs, independent of their appearance, remining so throughout the following starvation days (Figure 1A, Movie 1; Figure 1 – Figure supplement 1B). In order to quantify this drastic mobility change, we developed a procedure for automated quantification of the motion of LDs that were selectively labelled with the lipophilic dye Bodipy (Methods)(Listenberger & Brown 2007; Long et al. 2012). Simultaneous Phloxine B treatment enabled automatic exclusion of dead cells from analysis (Figure 1 – Figure supplement 2A; Methods) (Noda 2008). We were unable to use mean square displacement (MSD) as a measure to quantify LD mobility as phototoxicity prevented the prolonged imaging required. Instead, we determined the Pearson Correlation Coefficient (CC) for all pixels of each cell between two consecutive time points, a measure commonly used to quantify the co-localization of two fluorescent markers (Adler & Parmryd 2010; Dunn et al. 2011). We used it to quantify the “co-localization” of a fluorescent marker with itself at two time-points separated by 42 seconds (Methods). The maximal CC of 1 describes a completely static distribution of fluorescence intensity. Increased mobility of fluorescent structures will decrease the CC accordingly. In exponentially growing cells, we measured a CC of 0.56, which reflects the strong, seemingly erratic motion including occasional jumps of LDs that occurred throughout the cell volume (Figure 1B; Movie 1). After cells had entered a quiescent state at around SD2, LD mobility was reduced and LDs mainly wobbled in a given location without major translocations (Movie 1). Accordingly, the CC increased to 0.70 (Figure 1B). On SD3 there was little change with average median CCs of 0.68. On SD4 we measured a slight increase to 0.75. Only on SD5 did the average median CC start to increase significantly with an average median CC of 0.84. On SD6 it reached 0.966, consistent with the apparent, complete immobilization of LDs (Figure 1B). Thereafter, the average median CC remained high with 0.95 on SD7 and 0.95 on SD8. Remarkably, analysis of individual cell cultures showed that the mobility reduction of a given cell population was abrupt, occurring either on SD5 or SD6 (Figure 1C). On SD5 the average median CC of a given culture either remained closer to that measured for cultures on SD4, or it jumped to that measured for all SD6 cultures (Figure 1D). These results suggest that within a population, cells transit to the CF state in a coordinated fashion, within a relatively short period of time. Occasionally, we found a case where an entire cell culture did not transit to CF during SD5-SD8 (Figure 1 – Figure supplement 1C). Since such cell populations were rare and could be clearly distinguished, we excluded them from subsequent analysis of the CF state. Altogether, our measurements suggested the presence of two starvation phases, revealed by two abrupt decreases of LD mobility, which we termed early starvation and deep starvation. We termed this state where lipid droplets are immobilized throughout the cytoplasm of cells in deep starvation “cytoplasmic freezing” (CF).

**Figure 1.**
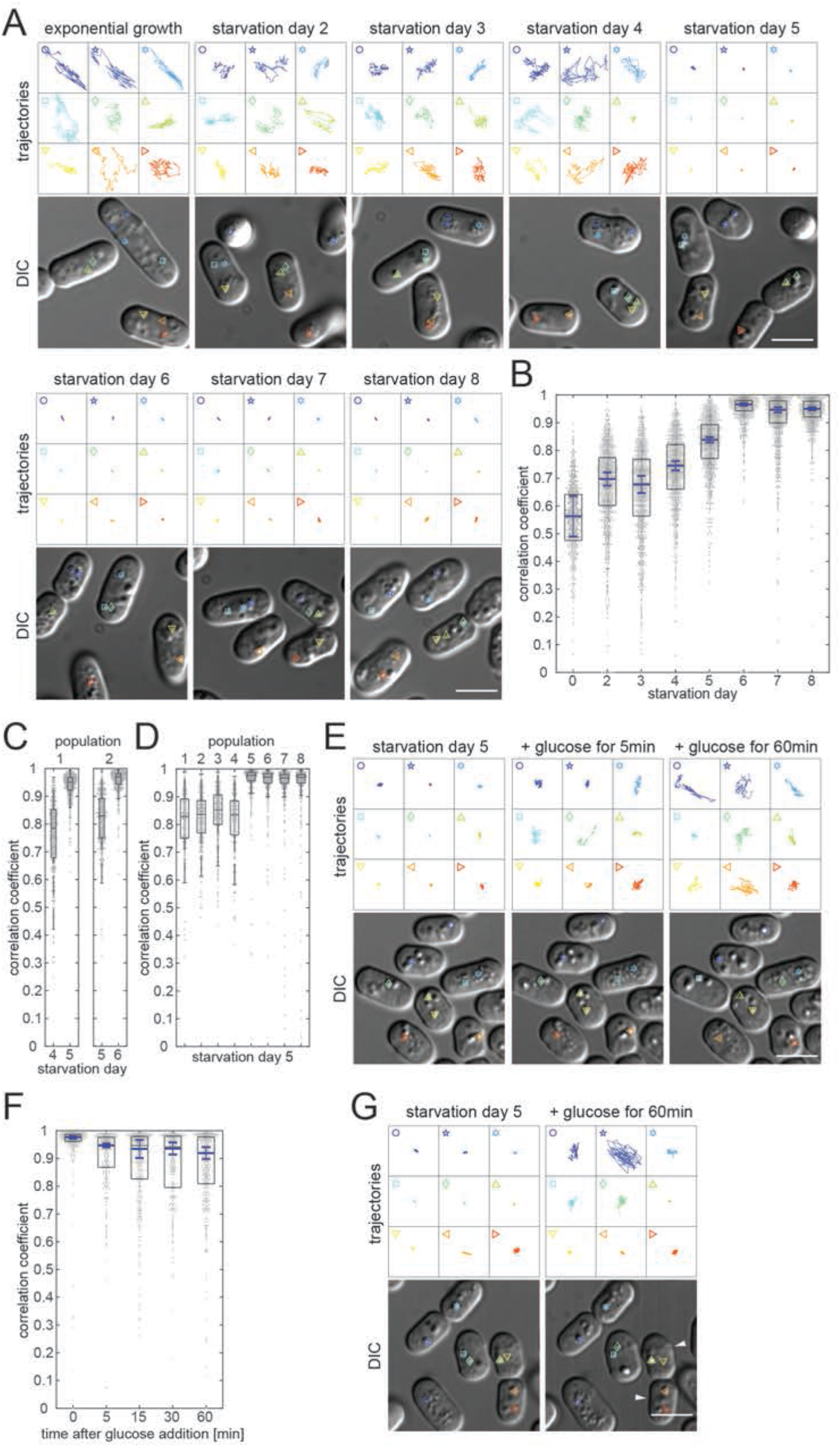
Motion arrest of lipid droplets in deep starvation. (**A**) Lipid droplet trajectories extracted from 25sec movies (4frames/sec, droplets depicted in lower DIC images) of cells during exponential growth (EG) and starvation days SD2-SD8. (**B**) Dot plots (one dot per cell) showing Pearson correlation coefficient-based (CC) quantification of Bodipy-labeled lipid droplet dynamics. Boxes represent the 25–75 percentile. The blue line represents the mean of 4 medians extracted from 4 independent cell populations (69 < n < 394 cells analyzed per experiment. Error bars show the 95% confidence interval. Note that none of the four cell populations entered CF on SD5. (**C**) Dot plot representing the CC of Bodipy-labeled lipid droplets in 2 independent cell populations (left and right panel) (325 < n < 372), revealing differing timing of transition to motion arrest (SD4–5 and SD5–6 respectively). Standard box plots are overlain. (**D**) Dot plot as in (C) representing the CC of Bodipy-labeled lipid droplets in 8 independent cell populations on SD5 (323 < n < 453). (**E**) Lipid droplet trajectories as in (A) of cells on SD5, immediately before and 5min or 60min after glucose addition. (**F**) Dot plot representation as in (B) showing quantification of lipid droplet dynamics at defined time points during starvation exit of cells from 3 independent cell populations (93 < n < 178 cells analyzed per experiment). (**G**) Lipid droplet trajectories as in (A) of cells on SD5 immediately before and 60min after glucose addition, showing lipid droplets remaining immobilized in two of the cells (white arrowheads). Scale bars in all panels: 5µm.

To investigate how cells regained LD mobility when exiting starvation, cells that had transitioned to CF on SD5 were supplied with fresh growth medium (Methods). Regained LD mobility was detectable within no more than 5 minutes, showing that the release occurs in a switch-like manner (Figure 1E; Movie 2). LD mobility increased over the following hour, which led to a broadening of the CC distribution (Figure 1F). Some cells remained in the CF state during this 1-hour observation period (Figure 1G, white arrows; Movie 2).

### CF differs from other solid-like cytoplasmic states

Two recent studies reported on a solidification of the cytoplasm in budding yeast cells that were acutely depleted of energy either by acute glucose depletion (AGD) or by “drug-induced energy depletion” (DED) (Joyner et al. 2016; Munder et al. 2016). Both studies showed some evidence for conservation in fission yeast. Since CF was very reminiscent of these other cellular states we directly compared acute energy depleted cells and cells in deep starvation (Methods). For DED cells, two protocols using 0.5h and 2h drug treatment were tested (Methods). Unlike in CF cells, LD motion was evident in AGD and DED cells when imaged with DIC microscopy (Figure 2A; Movie 3). Quantification with the Bodipy-labelling approach revealed an average median CC of 0.878 for AGD cells and of 0.900 for 0.5h DED cells, below the CC of for CF cells at 0.969 (Figure 2B). Interestingly, the average median CC of 0.966 for 2h DED cells did not differ much from that of CF cells, despite some clearly moving LDs. This could be due to the weaker Bodipy signal in CF cells, which might slightly reduce the precision of automated analysis.

**Figure 2.**
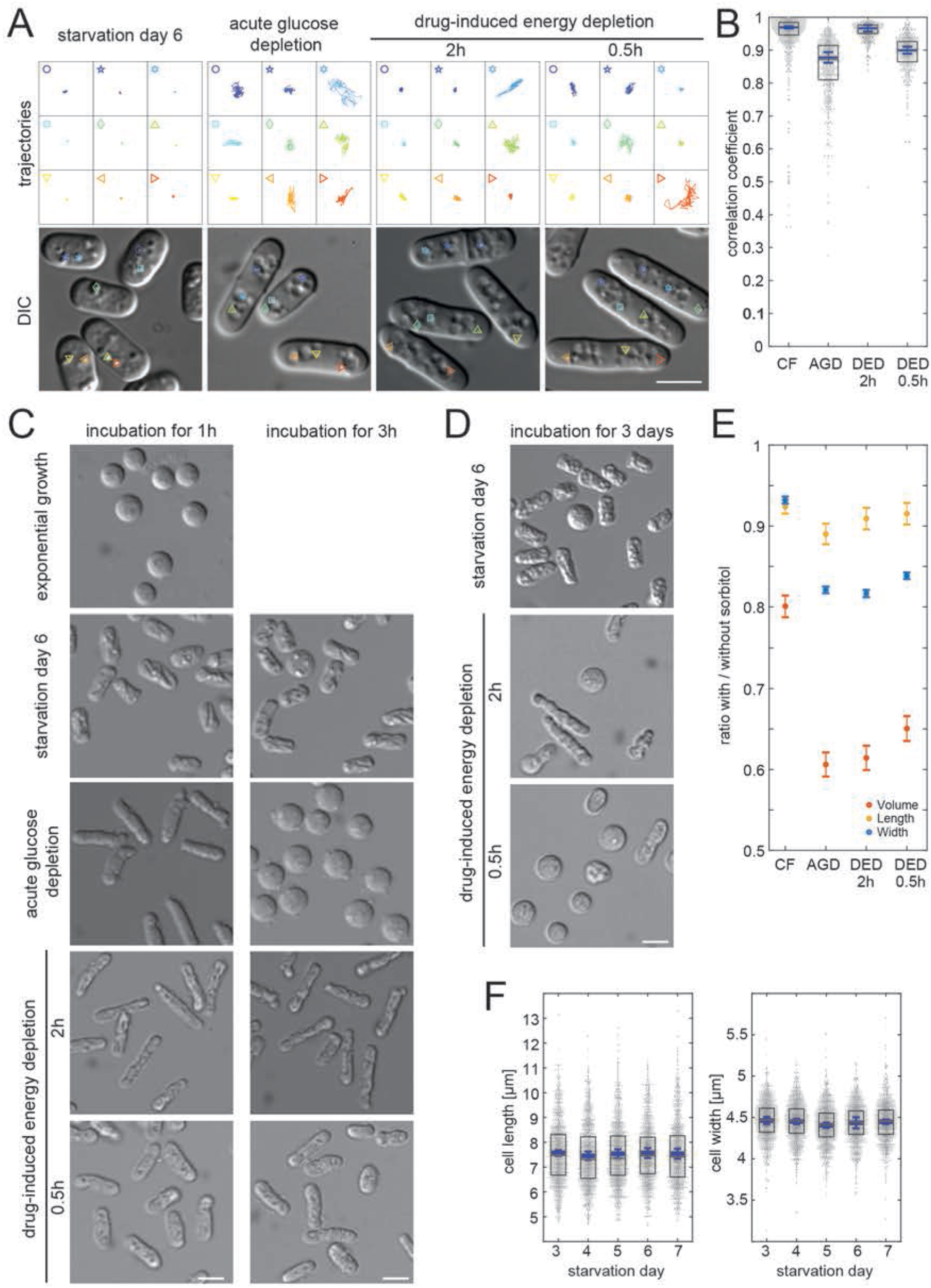
CF differs from other solid-like states and does not correlate with fluid loss. (**A**) Lipid droplet trajectories extracted from 25sec movies (4frames/sec, droplets depicted in lower DIC images) of CF cells on SD6, of AGD cells and of DED cells with 2h or 0.5h drug exposure. (**B**) Dot plots (one dot per cell) showing CC-based quantification of Bodipy-labeled lipid droplet dynamics in CF cells at SD6, in AGD cells and in DED cells with 2h or 0.5h drug exposure. Boxes represent the 25–75 percentile. The blue line represents the mean of the 3 medians extracted from 3 independent cell populations (73 < n < 312 cells analyzed per experiment). Error bars show the 95% confidence interval. (**C**) Protoplasts of cells incubated with cell wall digesting enzymes in 1.2M sorbitol containing buffer for 1h (left panel) and for 3h (right panel). (**D**) SD6 protoplasts from cells incubated as in (C) for an additional 3 days. (**E**) Ratios calculated from measured length (yellow), width (blue), or approximated volume (red) of cells placed in the respective standard culturing medium, or in 1.2M sorbitol containing buffer for 1h. To estimate the variance of the 3 experiments, bootstrapping was performed (Methods). Dots represent means, error bars show the 95% confidence interval of the bootstrapped mean ratios estimated with Gaussian error propagation. (**F**) Dot plots showing the cell length (left) and cell width (right) during starvation from SD3 to SD7. Boxes represent the 25–75 percentile. The blue line represents the mean of 3 means extracted from 3 independent cell populations (272 < n < 389 cells analyzed per experiment). Error bars show the 95% confidence interval. Measurements result from automated cell segmentation based on Phloxine B signal. Scale bars: 5µm.

To further test to what extent the cytoplasm of CF cells is solidified and how this compares to the state of cells with acute energy depletion, we generated protoplasts by digesting away the cell wall normally maintaining the cylindrical cell shape. When applying the standard protocol for cell wall digestion, exponentially growing cells exiting the cell wall shell immediately adopted a spherical shape (Figure 2C; Figure 2 – Figure supplement 1; Movie 4; Methods). AGD or DED cells however, remained cylindrical as previously shown (Figure 2C) (Joyner et al. 2016; Munder et al. 2016). Likewise, cell wall-depleted CF cells remained cylindrical, indicating a solid-like cytoplasm similar to AGD and DED cells (Figure 2C). In order to investigate the robustness of shape preservation, we extended the observation period of cells in the continued presence of the digestive enzymes (Methods). We found that the vast majority of CF SD6 protoplasts remained cylindrical even after three additional days of incubation (Figure 2D). In contrast, the initially cylindrical shape of AGD protoplasts was unstable as within 3 hours the vast majority had become spherical (Figure 2C). DED protoplasts showed an intermediate phenotype and with few exceptions remained cylindrical within the first three hours of incubation. After three days of incubation however, approximately 20% of the 2h-treated DED cells and 80% of the 0.5h-treated had turned spherical (Figure 2D). These results indicate that the content of AGD and DED cells - but not of CF cells - can still rearrange with differing kinetics and that cytoplasmic solidification in CF cells is more profound than in cells with acute energy depletion. It remains to be shown why for DED, a prolonged drug exposure increases cytoplasmic solidification.

A possible explanation for the highly robust solidification of CF cells is that their cytoplasm experiences massive macromolecular crowding. Since our protoplasting experiments were done using a standard hypertonic digesting mix (1.2M sorbitol), which dehydrates cells to relieve hydrostatic pressure and prevent cell rupturing, it is conceivable that the resulting increase in macromolecular crowding would contribute to the observed shape preservation (Methods) (Atilgan et al. 2015). To test to what extent intact AGD, DED, and CF cells can dehydrate, they were placed in a hypertonic buffer solution containing 1.2M Sorbitol. We measured the changes in cell length and width and used these values to estimate the cell volume assuming it to be a cylinder plus two half spheres representing the cell ends (Figure 2 – Figure supplement 2; Methods). In particular the reduction in cell width is a good indicator of fission yeast cell volume loss (Methods). We found that under these conditions AGD and DED cells shortened and considerably reduced their width (Figure 2E; Figure 2 – figure supplement 2). The resulting volume loss was 39.38% for AGD, 35.05% for 0.5h-DED and 38.64% for 2h-DED. CF cells in contrast, showed much less reduction in cell length, width, and volume (volume loss 19.94%) indicating that they comparingly lose little fluid under these conditions (Figure 2E). A possible explanation is that CF cells are already severely dehydrated and cannot shrink much further. To test this possibility, we measured the development of cell length and width with daily measurements from SD3-SD7. We found no change in cell length and cell width at around SD5 when cells enter the CF state (Figure 2F). Such a permanence of length shows that CF is inconsistent with a model where a fluid loss-based increase in macromolecular crowding underlies CF. Furthermore, electron microscopy data, showed no evidence for a more crowded cellular content in CF cells as compared to the wild type (Figure 6D).

Having shown that fluid loss is an unlikely cause for CF this does not exclude that it contributes to the shape preservation of cells lacking a cell wall. To test for this possibility, we removed the cell walls of AGD, DED, and CF cells in a lower concentration sorbitol solution (Methods). Under these conditions all exponentially growing cells rapidly lysed and no protoplasts formed, confirming that this solution is hypertonic (Figure 3A, Movie 5). Similarly, we observed lysis for all of the 0,5 and for a majority of the 2h-treated (Figure 3A). Approximately 15% of the 2h-treated DED cells remaining cylindrical. Time-lapse imaging revealed that these cells did not manage to fully exit the cell wall shell, whereby the parts that did exit changed shape suggesting a fluid content (Figure 3B, Movie 5). Of the AGD cells, approximately 50% immediately lysed as soon as cell walls were sufficiently perturbed (Figure 3A, Movie 5). The remaining cells adopted a spherical shape, squeezing out of the cell wall shell, in a similar manner to exponentially growing cells in the hypertonic digestion solution (Figure 3B, Movie 5, Movie 4). By contrast, most CF cells exited the cell wall shell as solid cylinders also in the hypotonic solution (Figure 3A and 3B, Movie 5). On average, 70% of these cells remained cylindrical throughout an additional incubation day (Figure 3C). Interestingly, in contrast to the shriveled and indented surfaces of CF protoplasts in hypertonic digestion solution, these protoplasts had a smooth surface suggesting that the peripheral part of CF cells can take up fluid, albeit not to an extent that the overall shape or volume are affected (compare Figure 2C and 3A). Taken together, these results indicate that unlike cells with acute energy depletion, CF cells stably fix the entire cellular content in a way that is insensitive to osmotic stress. This provides additional evidence against increased macromolecular crowding as basic cause of CF, while supporting a mechanism involving some kind of network that globally fixes cellular content.

**Figure 3.**
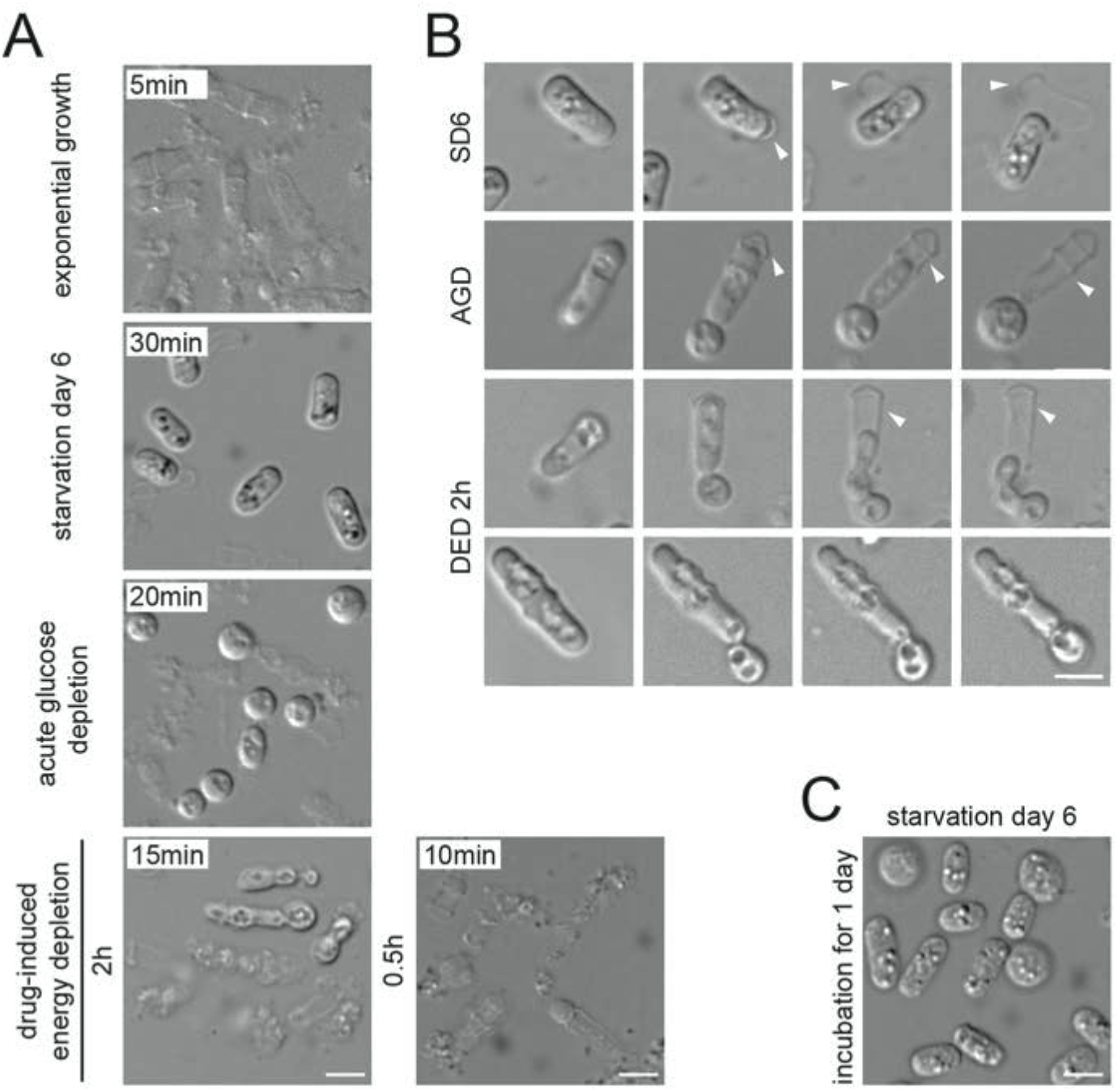
Protoplast generation under hypotonic conditions. (**A**) Protoplasts or cell remnants from cells incubated with cell wall digesting enzymes in 0.5M sorbitol containing buffer for the indicated times. (**B**) Live imaging of protoplast evasion from cell wall (white arrow) under 0.5M sorbitol conditions. Upper panel: CF cell, total time 6min; middle panel: AGD cell, total time 1.5 s; lower panels: DED 2h cells, total time 2min and 5min respectively. (**C**) CF protoplasts after 1 day of incubation with digesting enzymes in 0.5M sorbitol containing buffer. Scale bars: 5µm.

### Starved fission yeast lack extensive protein assemblies

In budding yeast, DED causes the homotypic assembly of a large number of proteins in a pH-dependent manner. Together, these assemblies were proposed to be responsible for the solidification of the cytoplasm. To check for a similar behavior of homologous proteins in fission yeast cells with DED, we followed eight of them tagged with GFP and/or mCherry when cells entered starvation in cells treated for 0.5h and 2h. Only for one of these we found cells in which the marker assembled into one or two spots following DED (Figure 4A; Figure 4 – Figure supplement 1A). However, only a subset of cells was affected. Another two proteins formed assemblies in exponentially growing cells when tagged with GFP(S65T), which did not change after DED (Figure 4B; Figure 4 – Figure supplement 1B). Curiously, when tagged with mCherry the protein did not form assemblies under any condition, suggesting that the assemblies of this protein were GFP(S65T)-dependent (Figure 4C; Figure 4 – Figure supplement 1C). All other proteins did not change their localization significantly (Figure 4D; Figure 4 – Figure supplement 1D). These results suggest that fission yeast cells do not react to DED by forming extensive macromolecular assemblies, unless by chance we selected unsuitable markers. Yet, our data suggest that GFP(S65T) is not suitable when testing for protein assemblies. It needs to be shown whether this is a general trend also for the budding yeast proteins, all of which were originally tagged with the GFP(S65T) variant (Huh et al. 2003).

**Figure 4.**
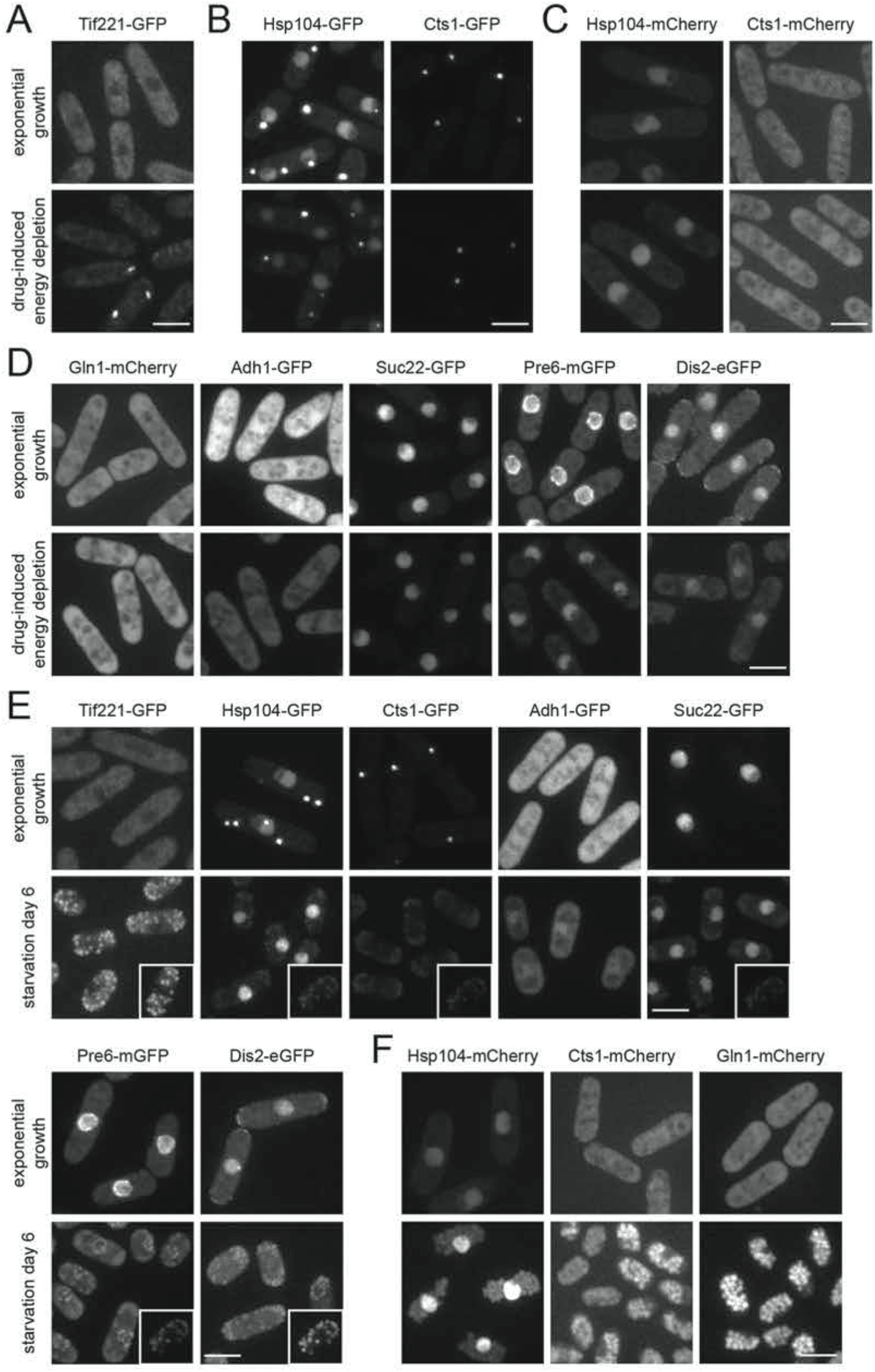
Fission yeast cells lack obvious macromolecular protein assemblies. (**A**)-(**D**): Images show fluorescence signal of the indicated fusion proteins in exponentially growing cells (upper panels) and DED cells (lower panels) incubated for 2 h prior to imaging. (**E**)-(**F**): Images show fluorescence signal of the indicated fusion proteins in exponentially growing cells (upper panels) and SD6 cells (lower panels). The unspecific signal portion can be estimated from comparison to the autofluorescence from a SD6 wild type cell without fluorescent tag with the same imaging and contrast settings (insets). Images are maximum intensity projections in all panels. All observations were confirmed in 2 independent experiments. Scale bars: 5µm.

While our data do not support the formation of protein assemblies in DED cells they do not exclude this possibility in CF cells. Therefore, we checked the behavior of all eight proteins in CF cells on SD6. When imaging SD6 of control cells not expressing any markers with a 488nm excitation laser and the GFP filter set, we noticed the gradual appearance of multiple autofluorescent particles (Figure 4 – Figure supplement 1E). By SD6, this autofluorescence had reached such a significant level that it had to be considered when imaging GFP-tagged proteins. The signal of most of the GFP-tagged proteins dropped below the autofluorescence level by SD6, suggesting that protein levels had significantly been reduced (Figure 4E). Importantly, no obvious aggregates formed and pre-existing aggregates disappeared as well. When proteins were tagged with mCherry, fluorescence mostly accumulated in large globular structures (Figure 4F). Co-expressing a vacuolar marker showed that the globular structures represented the vacuolar lumen (Figure 4 – Figure supplement 1F). The most likely explanation is that the mCherry tag accumulates in vacuoles after degradation of the attached proteins (Costantini et al. 2015). In one case, bright nuclear fluorescence accompanied the vacuolar labelling (Figure 4F). Supporting a complete absence of macromolecular assemblies in CF cells, electron microscopy pictures of CF cells showed no evidence of regions revealing protein aggregates as found in electron microscopy pictures of budding yeast cells (Figure 6D) (Petrovska et al. 2014). These results suggest that the levels of many proteins become significantly reduced in deep starvation and that extensive macromolecular protein assemblies are unlikely to be the basic cause of CF.

**Figure 5.**
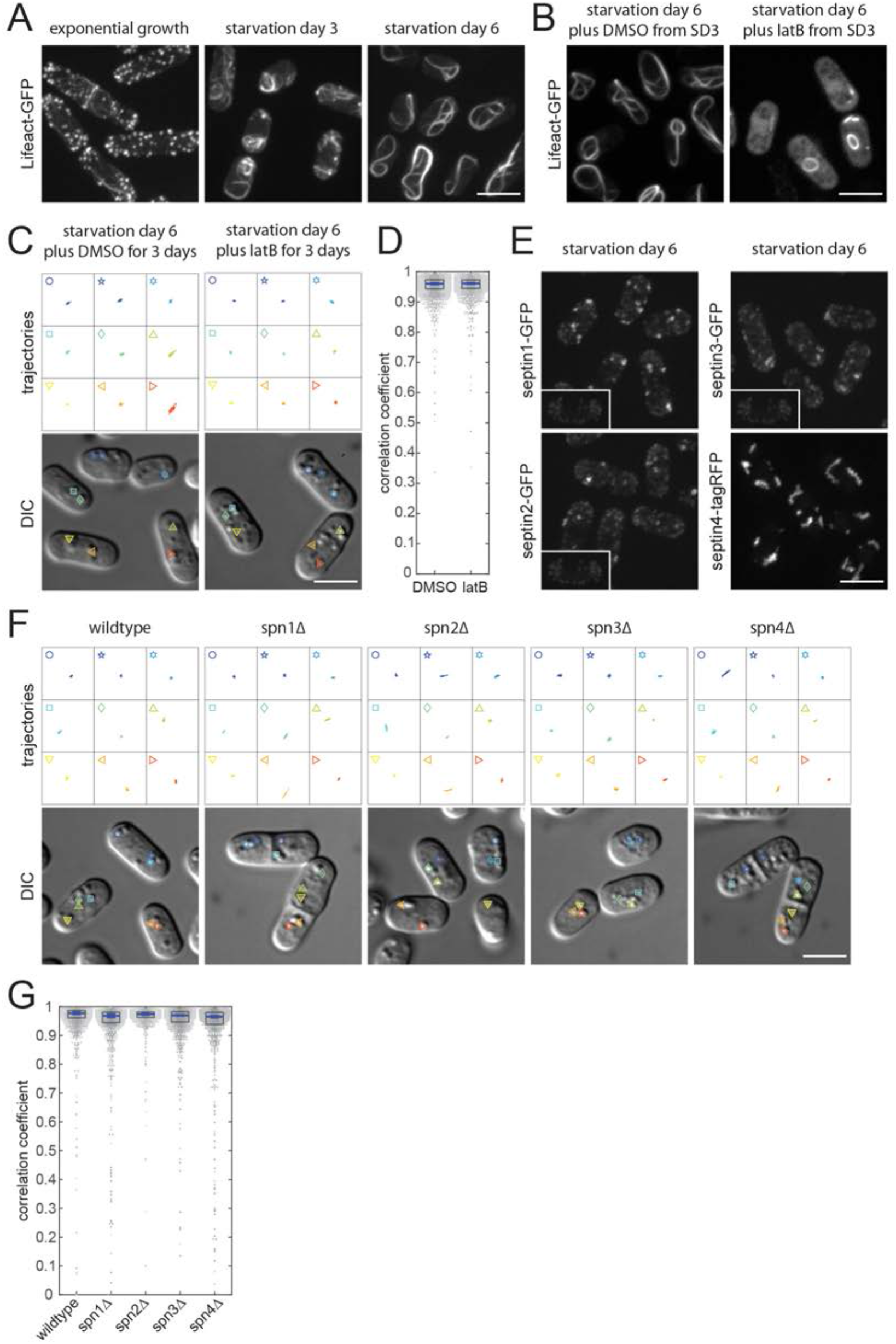
Cytoskeletal structures unlikely to be functionally relevant for CF. (**A**) Lifeact-GFP visualizing F-actin during starvation. (**B**) Lifeact-GFP from cells on SD6 that were incubated with LatB or DMSO from SD3 onwards. (**C**) Lipid droplet trajectories extracted from 25sec movies (4frames/sec, droplets depicted in lower DIC images) of wild type cells on SD6 incubated with DMSO or LatB from SD3 onwards. (**D**) Dot plots (one dot per cell) showing correlation coefficient-based (CC) quantification of Bodipy-labeled lipid droplet dynamics of wild type cells on SD6 from 3 independent cell populations incubated with DMSO or LatB from SD3 onwards. Boxes represent the 25–75 percentile. The blue line represents the mean of the 3 medians extracted from 3 independent cell populations (357 < n < 540 cells analyzed per experiment). Error bars show the 95% confidence interval. (**E**) Localization of Septin1–4 on SD6. The unspecific signal portion can be estimated from comparison to the autofluorescence from a SD6 wild type cell without fluorescent tag with the same imaging and contrast settings (insets) (**F**) Lipid droplet trajectories as in (C) of cells with spn1-spn4-deletions on SD6. (**G**) Dot plot representation as in (D) showing quantification of lipid droplet dynamics in cells with spn1-spn4-deletions on SD6 (265 < n < 476 cells analyzed per experiment). Fluorescence images represent maximum intensity projections. Scale bars: 5µm.

**Figure 6.**
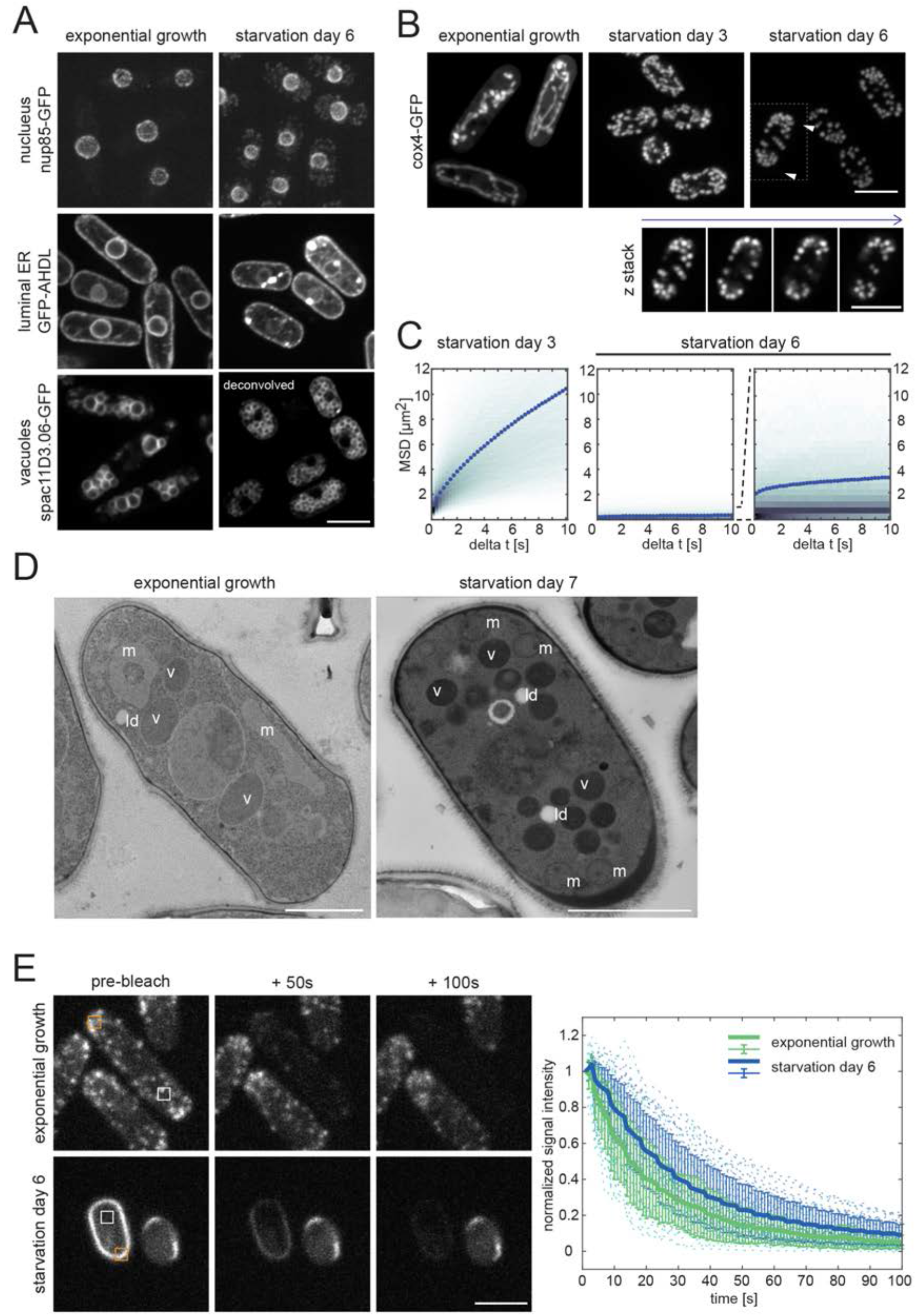
Subcellular architecture in deep starvation. (**A**) Nucleus (nup85-GFP), vacuoles (spac11D3.06-GFP), and ER (GFP-AHDL) at starvation day 6 (SD6) compared to exponential growth. The cytoplasmic dots in SD6 nup85-GFP cells represent autofluorescence rather than nup85-specific signal, see Figure 6 – Figure supplement 1. Images show single planes, deconvolved where indicated. (**B**) Mitochondria visualized by maximum intensity projections of cox4-GFP during starvation. The fragmented mitochondria on SD6 are often polarized (white arrows) and mostly cortical, as seen in the single planes of a z stack (lower panel; cell marked by the dotted square in upper panel) (**C**) Mean squared displacement (MSD) of spherical mitochondria particles at SD3 and SD6 (300 frames, 4fps, 6 independent experiments, 1766 < n < 4043 particles analyzed per experiment, 15 particles/cell on average). Plotted are color-coded histograms of each particle’s time-averaged MSD; Dotted lines show the ensemble- and time-averaged MSD’s. The second plot of SD6 shows a zoomed y-axis. In the first two panels y-axis numbers are x10^-3^, in the third panel x10^-4^. (**D**) Electron micrographs of freeze-substituted, plastic-embedded, and sectioned cells. The left panel shows a wild type cell in exponential growth and the right panel a spn3-GFP cell in the CF state on SD7. Starvation produces fragmented mitochondria, visible as small spheres marked (m in right panel), and remarkably different from regular, tube-like shapes (m in left panel). Vacuoles are marked (v) and low-density areas are labeled with (ld). Scale bar is 1µm. (**E**) Fluorescence loss in photobleaching (FLIP) experiments on Lifeact-GFP in exponential growth (upper panel) and on SD6 (lower panel). Images show maximum intensity projections of 3 planes. Repeated bleaching at the orange square every 5s. Plot shows bleach corrected and normalized mean signal intensity of white square. Thick lines: mean from 3 independent experiments (7 < n < 11 cells per experiment). Error bars: 95% CI of the mean; dotted lines: normalized signal of single cells. Scale bars: 1µm in (D) and 5µm in all other panels.

### CF occurs in the absence of the cytoskeleton

Having found no evidence of increased macromolecular crowding or protein assemblies causing the restriction of lipid droplet motion in CF cells, we investigated the possibility of structural fixation by a mesh-like protein network that might form throughout the cells. Tubulin and actin, which form the filaments of the cytoskeleton, are obvious candidates. Confirming previous results, we found that the multiple, dynamic microtubule bundles that are typical of growing cells remained active during SD2–4, but disappeared in deep starvation (Figure 5 – Figure supplement 1A; (Laporte et al. 2015; Makushok et al. 2016). If ever, only a single, very short microtubule stump remained that often could barely be discriminated from the autofluorescence signal. Since the electron microscopy images of CF cells did not show any evidence of cytoplasmic microtubules either (Figure 6D), we conclude that microtubules do not make a significant contribution to the CF state.

To test for a role of actin, we used cells expressing the F-actin marker Lifeact-GFP (Riedl et al. 2008; Huang et al. 2012). In exponentially growing cells, Lifeact-GFP labeled thin actin filament bundles that align parallel to the long cell axis and dynamic actin particles that concentrated at growing cell poles. On SD2-SD4, the actin particles disappeared except for a few remaining dynamic patches that were distributed throughout the cells. The thin interphase filaments were replaced by thicker, dynamic F-actin bundles (Figure 5A; Movie 6). These extended along the cell periphery and often curled around the cell ends or curled up inside the cells. On SD6, all actin patches had disappeared and the F-actin bundles had evolved into extremely prominent, very long F-actin bands (Figure 5A). These bands were completely immobile and extended along the cell circumference while curling around both cell ends to form a structure often reminiscent of a shoelace (Figure 5A; Movie 6). The strong signal of these actin bands indicates that it is likely to contain much of the actin protein pool normally present in cells. Yet, we could not exclude that the cells would additionally harbor a global Factin network, as single actin filaments cannot be detected by fluorescence imaging or electron microscopy as applied here. To prevent the formation of such filaments, we treated cells with LatrunculinB (LatB), a drug that interferes with F-actin polymerization (Methods) (Spector et al. 1983). Adding LatB to exponentially growing cells resulted in a fast depletion of all visible F-actin structures (Figure 5 – Figure supplement 1B). However, adding LatB to cells on SD6 did not affect the F-actin shoelace structures (Figure 5 – Figure supplement 1B). These results suggest that either these structures are no longer dynamic, or LatB cannot enter CF cells. Since this prevented drawings conclusions on the involvement of actin in CF, we applied the drug prior to the induction of CF on SD3 (Methods). This visibly decreased the dynamicity of the Factin structures but did not deplete them (Figure 5 – Figure supplement 1B). Interference with the formation of the shoelace-like actin cable, which was used to monitor the drug effect, was most efficient when incubating cells from SD3 onwards in the continued presence of the drug. On SD6 many cells ended up with a dispersed Lifeact-GFP signal although the marker still labelled short stumps and ring-like structures in a number of other cells (Figure 5B). LatB interference with the formation of the stable thick actin cables present in CF cells on SD6 suggests that the extended drug exposure would have blocked the formation of a putative global network of invisible, single actin filaments. Nevertheless, such cells showed CF on SD6 similar to DMSO-treated control cells (Figure 5C and 5D, Movie 7). We conclude that F-actin is not a major contributor to CF.

Septins represent another protein family that is known to assemble into filaments that could potentially form a global network in CF cells (Mostowy & Cossart 2012). The fission yeast genome has seven non-essential septin genes (*spn1*-*spn7*) (Longtine et al. 1996; Pan et al. 2007). Of these, spn5p-7p are exclusively expressed during meiosis (Abe & Shimoda 2000; Watanabe et al. 2001; Mata et al. 2002; Onishi et al. 2010). To ensure that they are not additionally expressed in deep starvation, we checked the signal of cells expressing GFP-tagged endogenous spn5p-7p on SD6. We found no signal above the autofluorescent signal and therefore carried on with the non-meiotic septins spn1p-spn4p (Figure 5 – Figure supplement 2B). When imaging these 4 septins tagged with GFP(S65T), we found that they all formed small clusters in deep starvation (Figure 5E). Next, we tested cells carrying a deletion of any of the four septins *spn1-spn4*, each of which affects proper formation of the septin ring in cytokinesis (Berlin et al. 2003; Tasto et al. 2003; An et al. 2004). On SD6 all septin deletion mutants showed CF that was indistinguishable from wild type (Figure 5F and 5G). These results make septins an unlikely candidate for mediating CF.

### Subcellular architecture changes in starvation

Since CF coincides with cytoskeleton rearrangements and appears to affect the entire cell, we initiated a general investigation on the organization of sub-cellular structures in deep starvation. For this, we used cells expressing various proteins tagged with GFP or mCherry that are known to mark cellular organelles and compared protein localization in exponentially growing cells with that of cells on SD6 (Table 1, strains used in this study). Only few markers maintained their localization. Of these, GFP-tagged nucleoporin nup85 and GFP-tagged AHDL – a marker of the nuclear membrane lumen and the endoplasmic reticulum respectively - showed a similar distribution in exponentially growing cells and in cells on SD6, despite some change in signal intensity. This indicates that these organelles do not reorganize much in deep starvation (Figure 6A). Similarly, the distribution of a GFP-tagged vacuolar marker did not change but the size of vacuoles appeared reduced on SD6 compared to exponentially growing cells (Figure 6A).

**Table 1:** Strains used in this study.

| Strains | Genotype | Source |
| --- | --- | --- |
| DB404 | h- Sec63-GFP::kanMX6 ura4-D18, leu1-32 | (Vjestica et al. 2008) |
| DB558 | h- wild-type | (Leupold 1950) |
| DB559 | h+ wild-type | (Leupold 1950) |
| DB933 | h- nup85-GFP::kanMX6 ade6-M216 leu1-32 | This study |
| DB2003 | h+ cnx1-HL-GFP::kanMX6 ura4-D18 leu1-32 ade6-M216 ** | This study |
| DB2057 | h- SV40p-GFP-atb2::LEU2 leu1-32 | (Pardo & Nurse 2005) |
| DB2400 | h+ anp1-GFP::ura4+ ade6-216 ura4-D18 leu1-32 | (Vjestica et al. 2008) |
| DB2401 | h+ sec72-GFP::ura4+ ade6-216 ura4-D18 leu1-32 | (Vjestica et al. 2008) |
| DB2402 | h+ anp1- mCherry::ura4+ ade6-216 leu1-32 ura4-D18 | (Vjestica et al. 2008) |
| DB2403 | h+ uch2-mCherry::ura4+ ade6-216 leu1-32 ura4-D18 | (Vjestica et al. 2008) |
| DB2404 | h- ost1::GFP-ura4+ ura4-D18 leu1-32 | (Vjestica et al. 2008) |
| DB2405 | h+ ost1-mCherry::ura4+ ade6-210 leu1-32 ura4-D18 | (Vjestica et al. 2008) |
| DB3287 | h- spn1Δ::kanMX6 | (Wu et al. 2010) |
| DB3293 | h- spn2Δ::ura4+ ura4-D18 | (An et al. 2004) |
| DB3297 | h+ spn2-GFP::kanMX6 * | (An et al. 2004) |
| DB3310 | h+ spn3-GFP::kanMX6 * | (An et al. 2004) |
| DB3324 | h- spn4-Δ1::kanMX6 * | (Wu et al. 2010) |
| DB3326 | h+ spn5-HL-GFP::kanMX6 | This study |
| DB3340 | h+ spn6-HL-GFP::kanMX6 | This study |
| DB3410 | h- leu1-32::pAct1-Lifeact-GFP::leu1+ * | (Huang et al. 2012) |
| DB3422 | h- spn1-GFP::kanMX6 * | (Wu et al. 2010) |
| DB3426 | h+ spn3-Δ::ura4+ ura4-D18 | (An et al. 2004) |
| DB3455 | h+ spn7-GFP::kanMX6 * | (Onishi et al. 2010) |
| DB3587 | h- spn4-tagRFP::kanMX6 | This study |
| DB3623 | h- Pbp1-GFP-AHDL::leu1+ ura4-D18 leu1-32 ade6 | (Zhang et al. 2010) |
| DB3624 | h- Pbp1-mCherry-AHDL::leu1+ | (Zhang et al. 2010) |
| DB3726 | h- cox4-GFP::leu2 leu1-32 * | (Fu et al. 2011) |
| DB3856 | h- hsp104-mCherry::kanMX6 (hsp104***) | (Coelho et al. 2013) |
| DB4192 | h- GMA12-GFP::ura4+ ura4-D18 * | (Wang et al. 2002) |
| DB4672 | h+ Pnmt1-TM-mCherry::leu+ | (Zhang et al. 2012) |
| DB5013 | h- atg8Δ::kanMX6 * | Bioneer M-4030H-U5, (Kim et al. 2010) |
| DB5018 | h- atg1Δ::kanMX6 * | Bioneer M-4030H-U5, (Kim et al. 2010) |
| DB5160 | h- cts1-HL-GFP::kanMX6 (ura7***) | This study |
| DB5162 | h- pre6-HL-mGFP::kanMX6 (pre6***) | This study |
| DB5209 | h+ suc22-GFP (rnr4***) | (Vejrup-Hansen et al. 2014) |
| DB5310 | h- gln1-mCherry::natR (gln1***) | (Coelho et al. 2014) |
| DB5315 | h+ dis2-NEGFP::ura4 ura4-D18 * (glc7***) | (Alvarez-Tabarés et al. 2007) |
| DB5320 | h+ adh1-GFP::kanMX6 * (adh2***) | (Sigova et al. 2004) |
| DB5380 | h+ cts1-mCherry::kanMX6 *(ura7***) | (Coelho et al. 2014) |
| DB5381 | h- hsp104-GFP::kanMX6 * (hsp104***) | (Coelho et al. 2013) |
| DB5470 | h- tif221-HL-GFP::kanMX6 (gcn3***) | This study |
| DB5730 | h+ nmt1::GFP-HL-spac11D3.06::kanMX gln1-mCherry::natR | This study |

\* auxotrophic alleles were eliminated by crossing to wild type (DB559) \*\* HL = “happy linker” (Methods) \*\*\* *S. cerevisiae* orthologs

Another marker that changed was cox4p (cox4p-GFP), which marks the tubular mitochondria spanning the entire length of exponentially growing cells (Figure 6B) (Yaffe et al. 2003). On SD3, cox4p-GFP labelled shorter mitochondrial tubes and multiple dynamic globular structures of uniform size, suggesting that the mitochondria were undergoing fission (Figure 6B; Movie 8). Consistently, on SD6 only globular mitochondria of uniform size were present, usually near the cell poles (Figure 6B). Similar to LDs, the motion of these globular mitochondria had completely arrested (Movie 8). Since cox4p-GFP fluorescence was sufficiently bright and stable, we could produce movie sequences that allowed determining the mitochondria MSD (Methods). We found an MSD close to zero in cells on SD6, confirming a nearly complete freezing of motion (Figure 6C).

For all other markers the signal marking the organelles vanished, when tagged with GFP, below the level of autofluorescence (Figure 6 – Figure supplement 1). As already described before, in cells expressing mCherry-tagged markers the mCherry signal ended up brightly staining the vacuoles on SD6 (Figure 4 – Figure supplement 1F). These data further support a general loss of protein diversity in CF, as was already suggested in our search for protein assemblies.

The global sub-cellular architecture of CF cells found by electron microscopy of plastic sections of cells collected on SD7 confirmed some of our observations with fluorescent organelle markers (Methods). First of all, we found no evidence for any kind of filamentous assemblies in CF cells (Figure 6D). In addition, confirming our fluorescence images, organelles such as LDs and vacuoles had the tendency to cluster around the nucleus in the cell center, while the almost spherical mitochondria with an approximate diameter of 250–350nm, were more prominent in polar regions near the cell periphery (compare Figure 1 – Figure supplement 1A with Figure 6B and 6D). In between these organelles, cells had had regions depleted of any distinct structure, except for electron dense spots typical of ribosomes. Altogether, our data suggest that the sub-cellular architecture of cells in deep starvation differs from that of growing cells and cells in early starvation.

### Immobilization of cytoplasmic components in CF is size-dependent

Having shown that fragmented mitochondria and LDs – both ranging in size between approximately 250µm and 350µm - were immobilized in CF cells, we wondered whether this also applies to components as small as individual proteins. The aforementioned Lifeact-GFP is a mainly globular, 29kDa protein of 4.5×2.5nm, binding to F-actin with a high turnover rate and otherwise seeming biochemically inert (Yang et al. 1996; Riedl et al. 2008). To test the mobility of the free, cytoplasmic Lifeact-GFP pool, we used the “fluorescence loss in photobleaching” (FLIP) method (Bancaud et al. 2010). FLIP involves repetitive bleaching of a defined, cellular sub-region. The kinetics of fluorescence loss in the remaining, unbleached cell regions provides a good measure of fluorescent particle motion into the bleached regions, thereby revealing information about the diffusive behavior of the particle. In CF cells on SD6, Lifeact-GFP depleted fast in the unbleached regions (Figure 6E). Notably, the strong signal on the stable actin cables also depleted fast. This reveals fast binding/unbinding of the fluorescent protein to actin as well as a rapid, diffusion throughout the cytoplasm that is comparable to the diffusion rate in exponentially growing cells (Figure 6E). We conclude that in CF cells, larger cellular components are fixed in place, while small molecules can diffuse almost as freely in CF cells as in exponentially growing cells. This is consistent with the presence of a global network structure with a certain mesh size in CF cells.

### Autophagy accelerates CF establishment

To further investigate the molecular nature of CF, we used our automated quantification of LD motion to screen for gene deletions that prevent CF in cells in deep starvation, using a library of 3400 fission yeast strains each carrying a deletion of a non-essential gene (Methods) (Kim et al. 2010). The strains were screened on SD8 since due to the presence of multiple auxotrophic mutations these strains entered deep starvation with a delay. Of the roughly 500 deletions that did not show CF in deep starvation, we noticed a clear accumulation of mutants affecting autophagy. Autophagy is an evolutionarily conserved mechanism used by cells to remove damaged organelles and to recycle cellular components (Nakatogawa et al. 2009). We further tested the contribution of autophagy to CF by taking a prolonged look at two strains each carrying a deletion of an essential component of the autophagy pathway, *atg1* (*atg1Δ*) or *atg8* (*atg8Δ*), in the absence of auxotrophic mutations. We found that both mutants entered the CF state but with a two- to three-day delay in comparison with the wild type (Figure 7A, 7B, Movie 9). Such *atg1Δ* and *atg8Δ* mutants robustly maintained a cylindrical shape after cell wall digesting in a hypotonic environment, again consistent with normal CF in these cells. These results show that rather than being a basic cause, autophagy helps cells to acquire a condition that allows them to induce CF. Thus, an unbiased gene screen identifies autophagy as a first cellular process required for CF.

**Figure 7.**
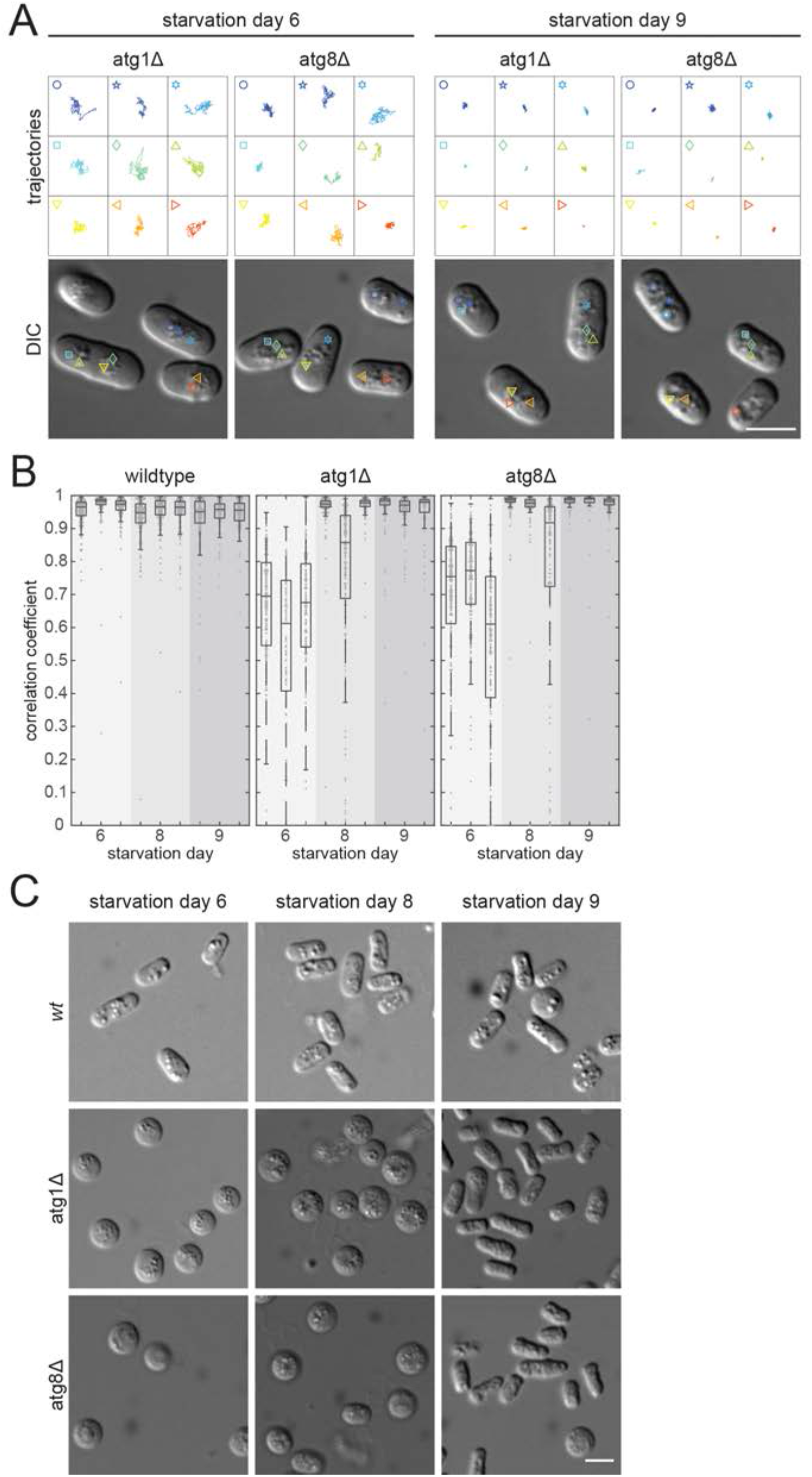
CF is delayed without autophagy. (**A**) Lipid droplet trajectories extracted from 25sec movies (4frames/sec, droplets depicted in lower DIC images) of atg1Δ and atg8Δ cells on starvation days SD6 (left) and SD9 (right). (**B**) Dot plots (one dot per cell) showing correlation coefficient-based (CC) quantification of Bodipy-labeled lipid droplet dynamics for wild type, atg1Δ, and atg8Δ cells separately for 3 independent cell populations (49 < n < 268 cells analyzed per experiment). Boxes indicate the 25–75 percentile. (**C**) Protoplasts of atg1Δ and atg8Δ at SD6 (left), SD8 (middle), and SD9 (right). Cell wall digestion for 4h in 0.5M sorbitol conditions. Scale bars: 5µm.

## Discussion

This work has uncovered a cytoplasmic solidification state, which we termed cytoplasmic freezing (CF) that provides the strongest solidification described for yeast cells to date. CF differs from the cytoplasm solidification states previously reported for budding yeast, acutely depleted from energy production in a number of key ways (Parry et al. 2014; Joyner et al. 2016; Munder et al. 2016). First, lipid droplet motion is completely abrogated in CF cells while it can be detected in cells with acute energy depletion. Second, we find ample evidence that fluid loss caused by a hypertonic medium, critically contributes to the previously reported cell shape preservation of AGD and DED protoplasts whereas CF protoplasts robustly maintain their cylindrical shape even in a hypotonic environment. Finally, neither fluorescence-nor electron microscopy revealed any evidence for the formation of widespread macromolecular protein assemblies as were proposed to mediate cytoplasm solidification in budding yeast cells with DED (Munder et al. 2016).

Altogether, our findings render increased macromolecular crowding a highly unlikely basic cause for CF. Instead they are compatible with a mechanism where CF cells contain a global network forming a grid with a defined mesh size that fixes larger cellular components but allows molecules of a size below the mesh size to freely move through the grid. At this point we don’t know anything about the nature of such a hypothetical network. We find no evidence for a role of the classical filament forming proteins of the cytoskeleton. A hint may come from the fact that even when placing CF cells in a hypertonic environment, we do not detect much fluid loss. This is not due to a complete insulation of cells from their environment, as they still take up dyes such as Bodipy and they still efficiently eliminate Phloxine B, our marker for automated detection of dead cells. Instead, such resistance to fluid loss would argue in favor of cells forming a hydrogel, which is typically resistant to dehydration (Tamai et al. 1996; Fullerton et al. 2006). A hydrogel could explain why standard electron microscopy pictures of CF cells do not reveal any network structures in the extensive free space between organelles. A hydrogel can form spontaneously via phase transition once the critical components have reached a certain concentration. This was proposed to occur during various processes, including Balbiani body formation, and it can involve proteins and/or RNAs (Brangwynne et al. 2009; Han et al. 2012; Kato et al. 2012; Patel et al. 2015; Boke et al. 2016; Jain & Vale 2017). In our case, such a phase transition may account for the tremendous synchrony with which CF occurs in a given cell population, either on SD5 or SD6 assuming that all cells in a population produce the critical components at a similar rate. It seems that quiescent cells need the 4–5 preceding days to become competent for switching into the CF state. In support of this, we find that blocking autophagy, a conserved mechanism for cell autonomous recycling of macromolecules, delays the onset of CF by 2 days. The most likely scenario is that cells use autophagy to cell autonomously provide energy and possibly the building blocks for specific components that mediate the CF state in the absence of external sources. This is remarkably similar to the process providing resistance to dehydration in tardigrades, where tardigrade cells need to express intrinsically unstructured proteins that eventually phase transit to a gel-like state fixing cellular content (Boothby et al. 2017).

Concerning the role of CF, it seems obvious that fixing cellular organelles and any other larger components in place provides a means to preserve the overall architecture of the cells. We cannot currently provide experimental evidence for a cytoprotective role of CF in deep starvation. However, it seems likely that, during an extended starvation period, the CF state will relieve cells from using vast amounts of energy to fight all the deleterious consequences of entropy. CF may also provide resistance to various stresses, without the need for metabolic activity. Unfortunately, our starvation conditions, which rely on standard fission yeast liquid culturing, are not optimal for investigating the role of CF in cell survival. The problem is that on around SD8, mortality rates massively increase. This is likely due to the fact that, under these conditions, cells enter starvation in their own, toxic waste. Thus, in order to perform such studies more favorable starvation conditions will first have to be established. Above all, we will need to find the factors that mediate CF. These will provide a means to interfere with CF establishment in order to test for advantageous features of this state. We expect the further analysis of the isolated deletion strains to reveal the critical components.

## Methods

### Yeast cell culturing

Cells were grown at 25°C in EMM2, supplemented with thiamine as required, and as described in (Moreno et al. 1991). For CF experiments, cells from a preculture in mid-exponential growth phase (EMM2 medium) were diluted to OD 0.02 (spectrophotometer Genesys 10S Vis, Thermo Fisher Scientific, Waltham, Massachusetts) in EMM, with 0.5% glucose (EMMLG) plus supplements as required. Cells were cultured in Erlenmeyer flasks on a shaker (New Brunswick scientific innova 4230; 220rpm) for up to 8 days. The culture volume did not exceed 1/10 of the total volume of the flask. All strains used are listed in Supplemental Table 1. For starvation exit, starved cells in a CF state were supplemented with fresh EMM2 (1:4) and incubated in an Erlenmeyer flask on the shaker at 25°C. AGD was done as described for budding yeast in (Joyner et al. 2016). DED was done as described in (Munder et al. 2016) (Antimycin A from Streptomyces sp., Merck, Darmstadt, Germany, A8674; 2-Deoxy-Dglucose, Merck, D8375). We performed all experiments with 0.5 and 2h drug incubation, as both incubation times were used in (Munder et al. 2016) for the various drugs tested.

### Protein tagging and constructs

Protein tagging was performed as described in (Bähler et al. 1998), using the primers listed in Supplemental Table 2, except that plasmids were equipped with a “happy linker” (HL) sequence between gene and fluorophore sequence where indicated as for EB1 in (Jankovics & Brunner 2006). mGFP(A206K) was generated from the routinely used GFP(S65T) by site-directed-mutagenesis at the dimer interface using forward (AGGTCGACGGATCCTTGGAG) and reverse (GATCTTTCGAAAGTTTAGATTGTGTGGACAGGTAATGGTTGTCTGG) primers, and insertion into the original plasmid using restriction enzymes BamHI / BstBI (Snapp 2003).

**Table 2:**
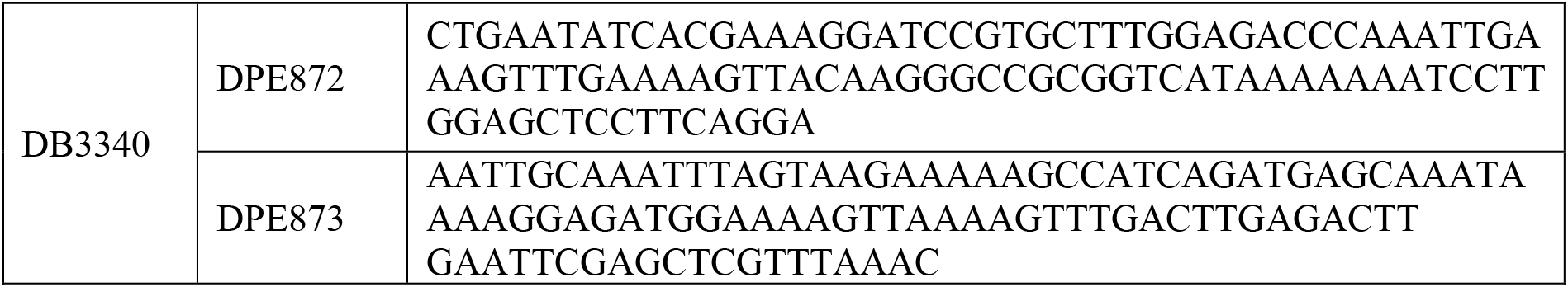
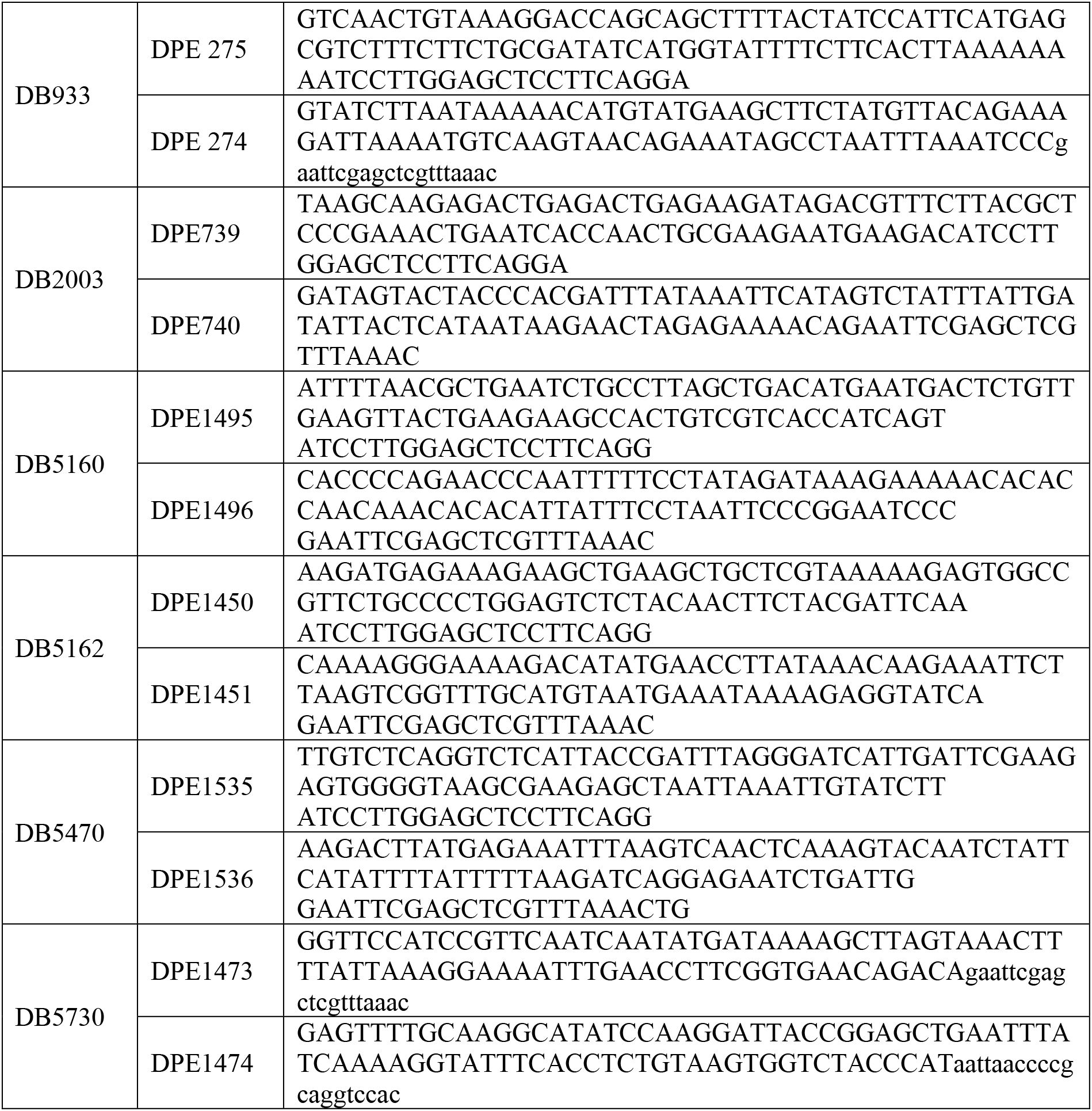
Primers used in this study.

### Microscopy

Live imaging was performed at room temperature (RT) on a spinning disc microscope (Zeiss Axio Observer Z1, Yokogawa CSU-X) using 63x and 100x NA 1.4 oil plan apo objectives, Andor iQ2.9 software, Andor Neo sCMOS and iXon3 EMMCCD cameras (Andor Technology, Belfast, UK) using 488nm and 561nm laser excitation and 525/50 BP and 568 LP emission filter sets. Z stacks with 0.5µm steps were acquired for standard fluorescence unless stated otherwise.

DIC imaging of starved cells was performed on poly-L-lysine (2mg/ml, Merck, P1399) coated glass bottom dishes (Bioswisstec, Schaffhausen, Switzerland; 5160). In the dishes, we constructed a chamber by adding a cover slip on top of 3 parafilm strips acting as spacers. The dishes were then placed for 2s on a heating plate at 100ºC, which partially melted the strips such that they glued the coverslip to the dish. The thereby formed chamber was filled with cell culture (∼30µl) by pipetting into the gap between the parafilm strips. To extract LD trajectories from DIC movies, cells were centrifuged at low speed (300rpm, Multifuge 1S-R, Thermo Fisher Scientific) and imaged immediately (5–15 min after mounting) as prolonged residence in the chamber caused artefacts. For each movie, 100 image frames were acquired at 4fps.

LD trajectories of cells exiting starvation (Figure 1E) were extracted from DIC movies using lectin-coated glass bottom dishes (griffonia (bandeiraea) simplicifolia lectin1; Vector laboratories, Burlingame, California; L-1100). A first movie was taken from cells in EMM without glucose (EMM0G) immediately before glucose addition. After 2% glucose addition, movies were taken for up to 60min.

Live cell imaging for CC quantification was done on lectin coated glass bottom 8-well (ibidi, Martinsried, Germany; 80827) or 10-well slides (Greiner Bio-One, Kremsmünster, Austria; 543079) after centrifugation at 1000rpm. 1mg/ml Bodipy (BODIPY 493/503; Thermo Fisher Scientific; D3922) was dissolved in DMSO and used at a final concentration of 4µg/ml in EMM2 or EMM0G for exponential or starved cells respectively. Phloxine B (Merck, P4030) was dissolved in water to 5mg/ml, diluted to 100µg/ml in water, and used at a final concentration of 10µg/ml. When imaging, we first acquired a single focal plane image of red fluorescent Phloxine B followed by 3 single focal plane images of green fluorescent Bodipy with a time interval of 42s.

The movies for cox4-GFP particle tracking were made of 300 frames taken at 4fps in chambers as for DIC imaging.

Protoplasts in high sorbitol were imaged with DIC on lectin coated glass bottom 10-well slides after centrifugation at 1000 rpm. Protoplasts in low sorbitol were transferred to a lectin coated imaging chamber (see above), sealed off with VALAP (Vaseline, Lanoline, Parafilm; 1:1:1) to prevent dehydration, and imaged immediately.

### Cell wall digestion

Protoplasts in high sorbitol were generated by enzymatic digestion of cell wall, with slight alterations to a previously published protocol (Kelly & Nurse 2011). To form protoplasts, cells were incubated at 25°C with 5mg/ml Zymolyase 20-T (MP Biomedicals) plus 5mg/ml lysing enzymes from Trichoderma Harzianum (Merck; L1412) in 500uL E-buffer +1.2M sorbitol in a 2mL Eppendorf tube for 1h on a rotor at 25°C unless stated otherwise.

Protoplasts in low sorbitol were generated by washing cells in E-buffer + 0.5M sorbitol, centrifuged at minimal speed for 5min, and resuspended in 50uL E-buffer + 0.5M sorbitol plus cell wall digesting enzymes.

For DED, protoplasts were generated in continued presence of 20mM 2-deoxy glucose and 10mM antimycin A in E-buffer as described in (Munder et al. 2016).

### FLIP acquisition and analysis

Fluorescence loss in photobleaching (FLIP) experiments were performed on cells mounted to an imaging chamber sealed with VALAP (see above). Imaging was done at RT on a spinning disc microscope (Nikon Eclipse Ti, VisiScope system, Yokogawa W1) using a 60x water objective, VisiView software, and an Andor EMCCD camera (iXon Ultra 888 back illuminated). A z-stack of 3 planes (1µm step size) was acquired every second for 100sec while a small region with 1.12×1.12µm size near one cell pole was bleached every 5sec. The mean fluorescence intensity loss of a reference region at the opposite pole was then extracted using Fiji. The analysis was done using Matlab, as described in (Bancaud et al. 2010). The signal was normalized to the last pre-bleach time point. For each condition, 30 cells were analyzed - 10 each in three independent experiments.

### Electron microscopy

Cells were high pressure frozen in solution (reviewed in (McDonald et al. 2010)) using a Wohlwend Compact-2 high-pressure freezer (Martin Wohlwend AG, Sennwald Switzerland). *S. pombe* samples destined for plastic section microtomy were freeze-substituted in 0.1% glutaraldehyde and 1% uranyl acetate in acetone for 48hrs and warmed from –90°C to –50°C in 8hrs (5°C per hour). Cells were then washed by acetone for 3 times and infiltrated in HM20 solution (25%, 33%, 50%, 67%, 75%, 100% in acetone) (Lowicryl HM20 Embedding Kit, Electron Microscopy Science, Hatfield, PA) over 5 days using Leica EMAFS (Leica, Vienna, Austria). Samples were then polymerized to blocks under Leica EMAFS UV light unit for 72hrs.

Plastic blocks were cut into ribbons of 80 (for single projection images) – 250 nm thick plastic sections (for tomographic reconstructions), depending on the questions asked, by Leica Ultracut microtome (Leica Inc., Vienna, Austria) using Diatome Ultra 45° (Diatome AG, Biel, Switzerland). Ribbons were collected on formvar-coated Cu-Rn grids (Electron Microscopy Science, Hatfield, PA) or Carbon Film Finder grids (Electron Microscopy Science, Hatfield, PA), immuno-labeled (optional), stained by uranyl acetate (2% uranyl acetate in 70% methanol) for ∼4min and Reynold’s lead citrate for ∼2min (the staining time was adjusted based on the thickness of the sections).

Individual pictures of plastic sections, mostly used as a control, were acquired with a FEI Philips CM100 TEM and AMT 2Kx2K bottom-mount digital camera.

### LatB treatment

LatB (Latrunculin B, Latrunculia magnifica, Merck; 428020) was added from a stock solution of 10mM in DMSO to a final concentration of 100µM (1% DMSO). Control cultures were treated with 1% DMSO. For the short-term effect of this LatB concentration, the stock solution was diluted in EMM2 for exp. cells and EMM0G for SD3 and SD6 cells. For 3-day LatB incubations, starved cultures were split in half at SD3. One half was supplemented with LatB, the other with DMSO to serve as control. Both cultures were incubated at 25°C on the shaker for another 3 days.

### Image Analysis

Routine image processing was done using Fiji/ImageJ. Deconvolution was done using Huygens software (Scientific Volume Imaging) on image stacks acquired using Nyquist criteria. Plots were made using Matlab (MathWorks).

#### LD trajectories from DIC movies

The DIC movies (100frames, 4fps) were stack registered (Fiji plugin “StackReg”). Lipid droplets were tracked with the Fiji plugin “Manual Tracking with TrackMate” (settings for semi-automated tracking: Quality threshold: 0.2, Distance tolerance: 0.1, Max nFrames: 0). The lipid droplets were manually seeded in the first time-frame, and the trajectory was considered if the particle could be tracked for more than 95/100 frames. The manually seeded first trajectory point was excluded from the final trajectory, such that all LD positions were automatically detected. The trajectories were plotted using Matlab.

#### Mean square displacement of mitochondria

Cox4-GFP labeled mitochondria were tracked using the Mosaic particle tracking plugin in Fiji (Mosaic Toolsuite, (Sbalzarini & Koumoutsakos 2005); Settings: radius: 4, cutoff: 0, per/abs: 2, link: 3, displacement: 2). The subsequent analysis was done in Matlab (Mathworks). Only trajectories longer than 160 frames were considered, and the MSD up to a time lag of 40 frames was computed. The time-averaged MSD for each particle is plotted as a color-coded histogram, and additionally the ensemble-averaged mean is shown.

#### CC quantification of lipid droplet motion

Cell segmentation: The Phloxine B signal was log transformed and background subtracted in Fiji (Mosaic ToolSuite). Using pixel classification in ilastik the cell’s inside was separated from the outline and the background. Phloxine B-filled, and therefore dead cells, were marked as a separate class and excluded from subsequent analysis. Cell insides were segmented in Cellprofiler and used as seeds to segment the living cells (Figure 1 – figure supplement 2A). Subsequently, the 3 Bodipy images, taken at a time interval of 42s (t1-t3), were stack registered in Fiji (StackReg). We included additional procedures to account for high variability of Bodipy signal intensity amongst individual cells in deep starvation as well as significant differences of entire populations between different starvation days, with particularly low signal at SD2 and SD3 (Figure 1 – figure supplement 2B). In addition, for some unknown reason, Bodipy signal intensity gradually increased during imaging, with a major increase between t1 and t2 (Figure 1 – figure supplement 2C). As a result, the software generally detected LDs most efficiently at t2 and t3, which is why we use these timepoints to determine the CC. Because absolute signal intensities are irrelevant when extracting LD dynamics, we equalized fluorescence on our images by performing a log transform and background subtraction (Mosaic ToolSuite). Pixel classification in Ilastik further improved equalization. This generated pseudo images in which pixel values represent the probability to belong to a lipid droplet instead of the actual fluorescence intensities (Figure 1 – figure supplement 2D). From these, we computed the Pearson correlation coefficient (CC) for all pixels of individual cells between t2 and t3, using Cellprofiler. We present individual CC values as dot plots using the function plotSpread from the MathWorks File Exchange (https://ch.mathworks.com/matlabcentral/fileexchange/37105-plot-spread-points--beeswarm-plot-). We overlay a boxplot indicating the median and the 25^th^ to 75^th^ percentile of the values. Where several experiments were pooled, the mean of the medians of the single experiments was plotted in blue. As a measure for the variance between individual experiments, blue error bars indicate the 95% confidence interval (95% CI) of the medians. This variance does not describe the variance of the individual medians and might thus underestimate the true variance of the median.

#### Cell size measurements

For comparison of cell size in respective standard culturing and 1.2M sorbitol containing medium (Figure 2 E, Figure 2 – figure supplement 1 C), cell length and width were measured manually from DIC images using Fiji. Subsequently, we extracted the means of cell length and cell width measured from 3 independent experiments each, for cells culturing in standard and in high sorbitol medium. To estimate the variance (*dL^2^, dW^2^*) of the 3 experiments, bootstrapping was performed (999x resampling of each individual experiment), leading to 999 new means from bootstrapped samples. The mean cell volume *V* was approximated as a cylinder plus a ball from the mean of the bootstrapped means of measured length and width (*L, W*) as follows 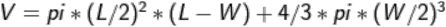. The variance of the cell volume (*dV^2^*) was estimated by Gaussian error propagation with 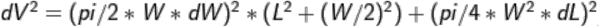. To define the amount of cell shrinkage after transfer to 1.2M sorbitol containing medium, we divided the mean of the bootstrapped means of cells in 1.2M sorbitol containing medium (*S*) by the mean of cells in respective standard culturing medium (*E*) for length, width, and approximated volume. The variance (*dR^2^*) of the ratio 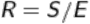 was estimated by Gaussian error propagation with 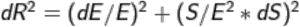. The mean and 95% CI of these normally distributed fractions are shown in Figure 2E.

Measurements of cell length and width using Phloxine B signal was done in an automated fashion as described above.

---

**Figure 1 – figure supplement 1.**
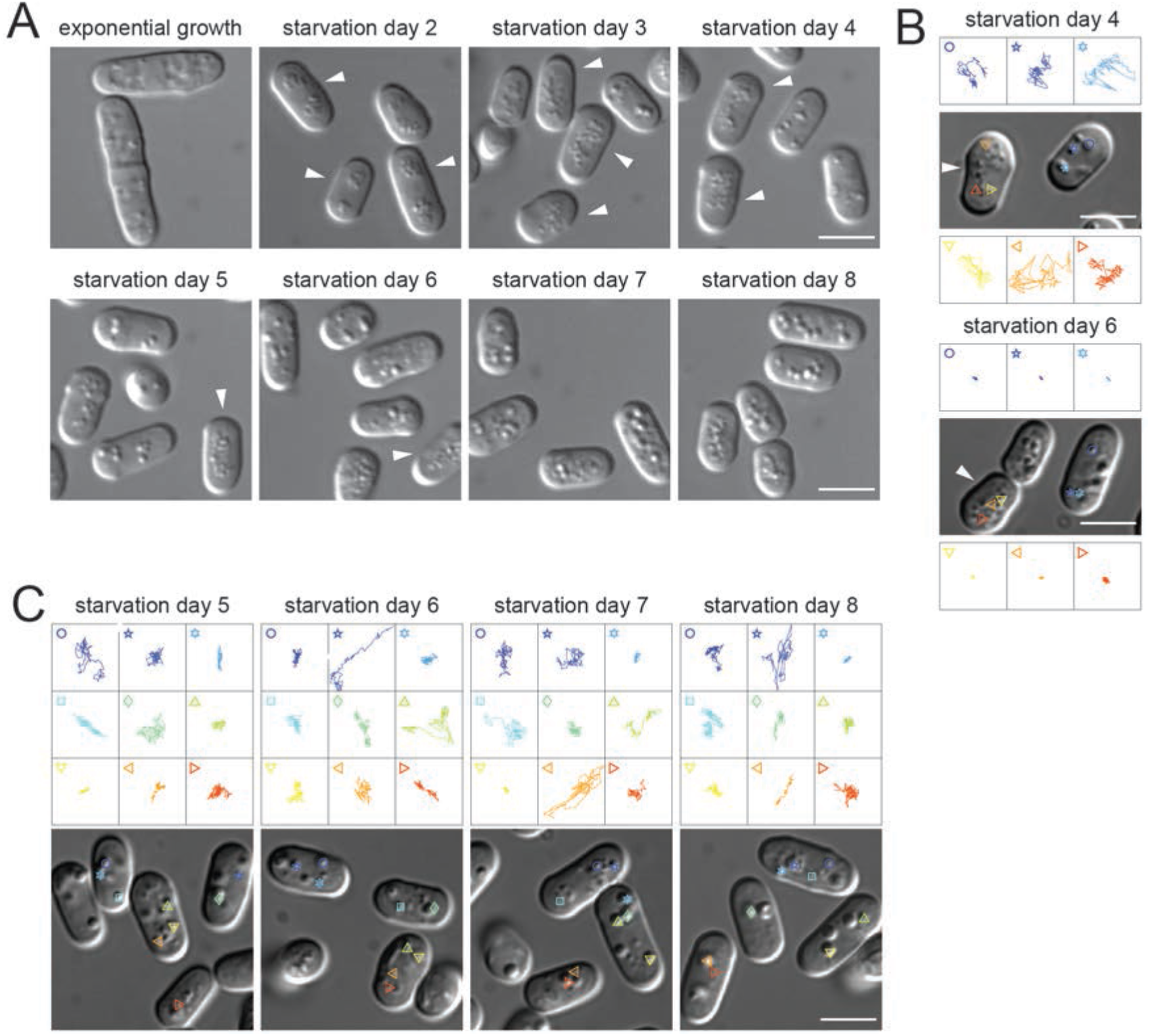
Lipid droplet morphology and dynamics in starvation. (**A**) Lipid droplet composition changes from evenly distributed droplets in exponentially growing cells to grape-like lipid droplets (white arrows) in cells on SD2. From SD3 to SD5 cells increasingly contain fewer but bigger lipid droplets. (**B**) Lipid droplet trajectories as in Figure 1A, comparing cells containing grape-like lipid droplets (white arrows; orange trajectories) and cells containing bigger lipid droplets (blue trajectories). Upper panel shows cells on SD4, the lower panel on SD6. (**C**) Lipid droplet trajectories as in Figure 1A, of a cell population in which no motion arrest occurred up to SD8. Scale bars in all panels: 5µm.

**Figure 1 – figure supplement 2.** Determining the correlation coefficient of lipid droplet dynamics.

(**A**) Images show Phloxine B-labeled cells (upper panel) as used to segment cells. Segmented cell outlines are shown in green (lower panel). Dead cells internalized Phloxine B (white arrow) and were excluded from segmentation and subsequent analysis. (**B**) The upper panels show the Bodipy signal in the cells of the DIC images in the lower panels, during exponential growth, on SD3 and on SD6. White arrows depict individual cells with an extreme difference in Bodipy signal intensity. (**C**) Bodipy-labeled cells on SD3 imaged at 3 consecutive time points (t1-t3), separated by 42s. Note that contrast settings for the same pictures differ in (B) and (D). (**D**) Same cells as in (B) showing the Bodipy image pixels in the upper panels and in the lower panels the respective pseudoimages resulting from classification with Ilastik. The pseudoimages show the probability of pixels to belong to lipid droplets as was then used to compute the CC between two imaged time points. Scale bars: 5µm.

**Figure 2 – figure supplement 1.**
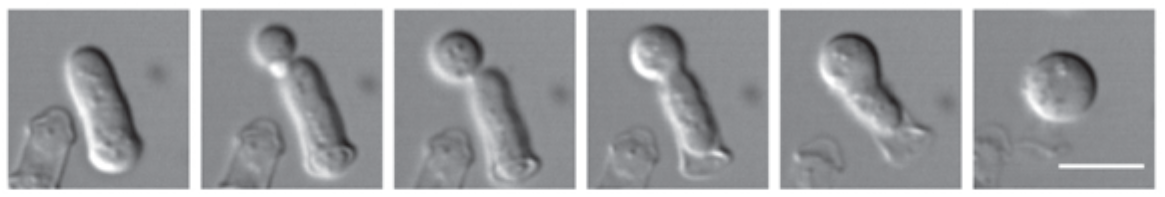
Cell wall evasion of an exponentially growing cell. Cell wall digestion in 1.2M sorbitol. Total duration 5min. Scale bar: 5µm.

**Figure 2 – figure supplement 2.**
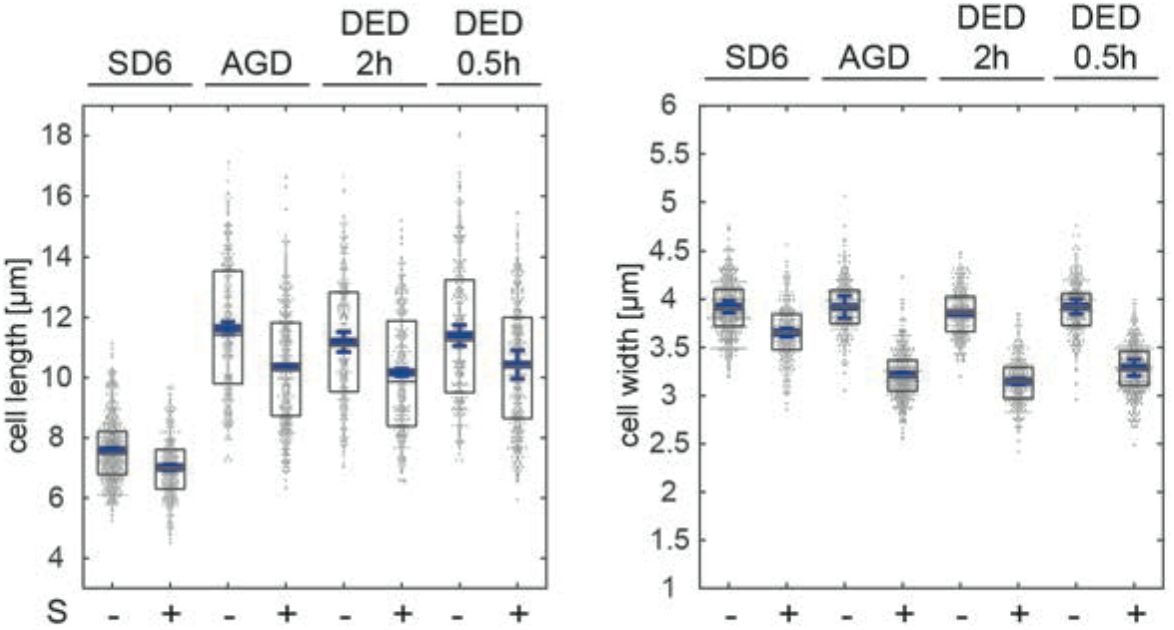
Cell size in normal medium and 1.2M sorbitol buffer. Cell length and width of CF, AGD, and DED (2h and 0.5h) cells in standard culturing medium (-S) or in 1.2M sorbitol containing buffer (+S) from 3 independent cell populations each, measured manually from DIC images. The blue line represents the mean of 3 experimental means (103 < n < 269 cells analyzed per experiment). Error bars show the 95% confidence interval.

**Figure 4 – figure supplement 1.**
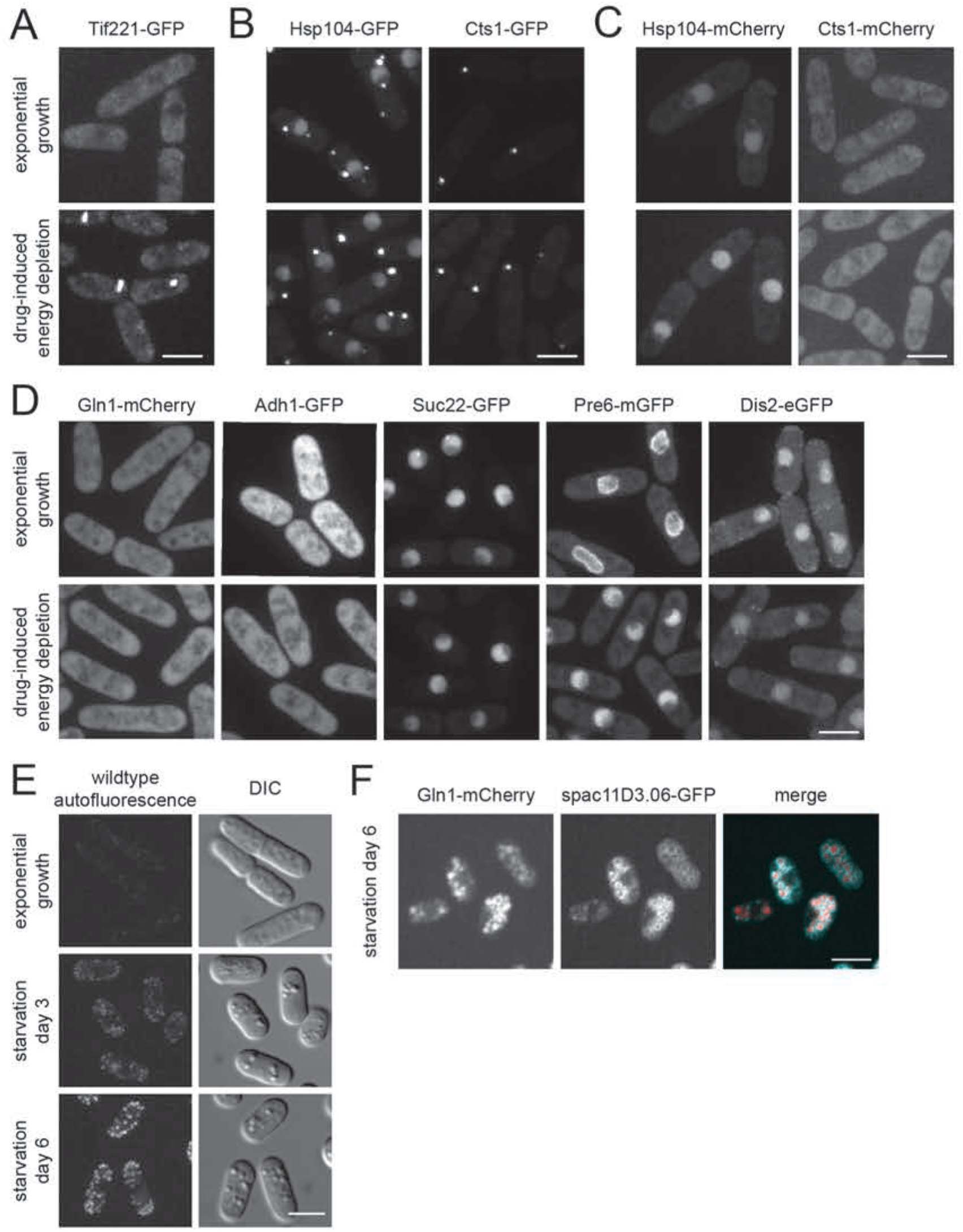
Fission yeast cells lack obvious macromolecular protein assemblies. (**A-D**) Images show fluorescence signal of the indicated fusion proteins in exponentially growing cells (upper panels) and DED cells (lower panels) incubated for 0.5 h prior to imaging. (**E**) Autofluorescence (excitation 488nm, emission 525/50) of wild type cells in exponential growth, on SD3 and SD6. Images show maximum intensity projections. (**F**) Single focal plane images showing double labeling of Gln1-mCherry and the vacuolar marker spac11D3.06-GFP. Scale bars: 5µm.

**Figure 5 – figure supplement 1.**
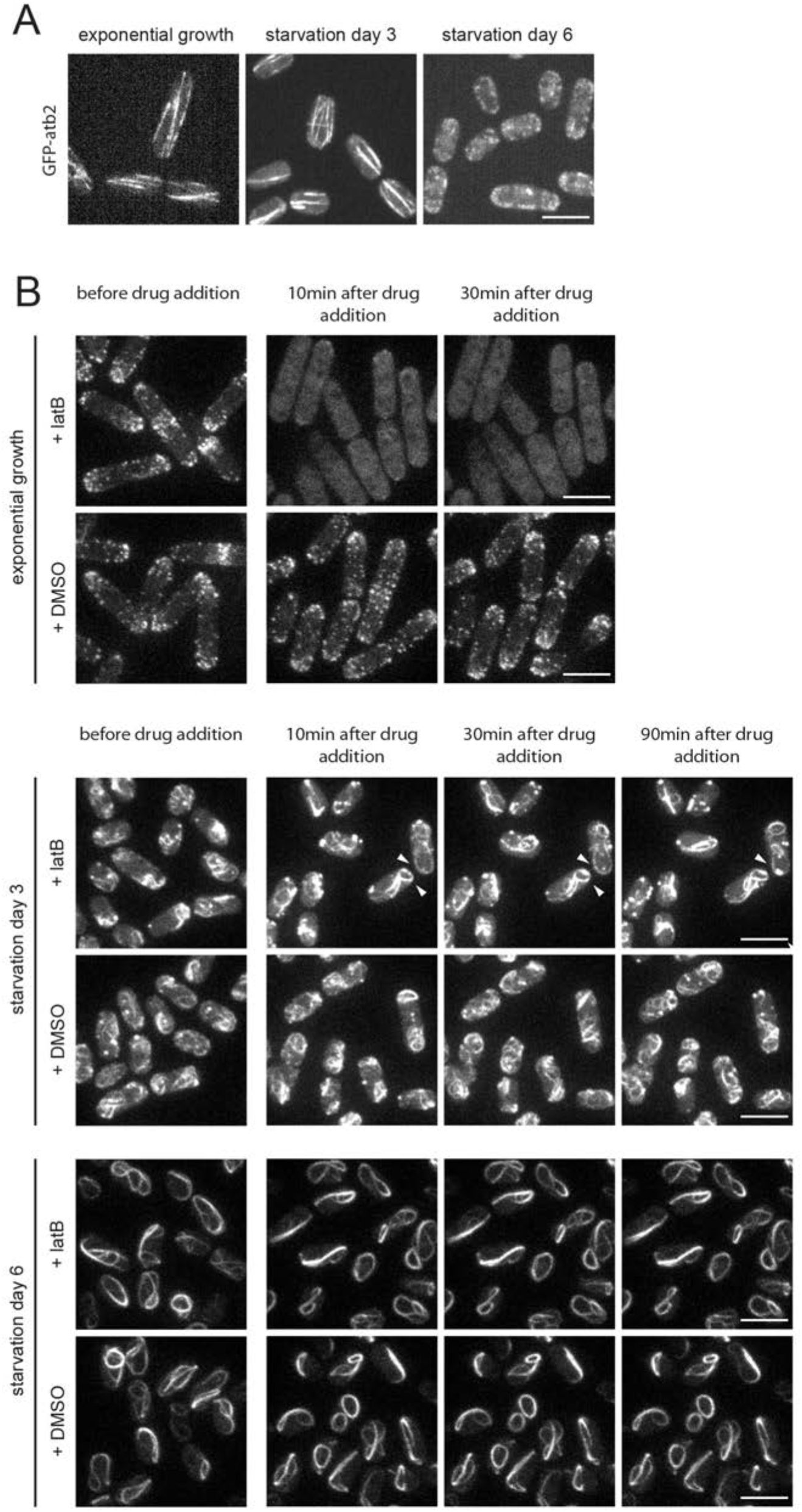
Microtubules during starvation and acute effect of LatB. (**A**) Microtubules visualized by GFP-atb2 during starvation. (**B**) Effect of 100µM LatB/DMSO on F-actin in exponential cells (upper panel), SD3 cells (middle panel; white arrows indicate cells with reduced F-actin dynamics), and SD6 cells (lower panel). Images represent maximum intensity projections. Scale bars: 5µm.

**Figure 5 – figure supplement 2.**
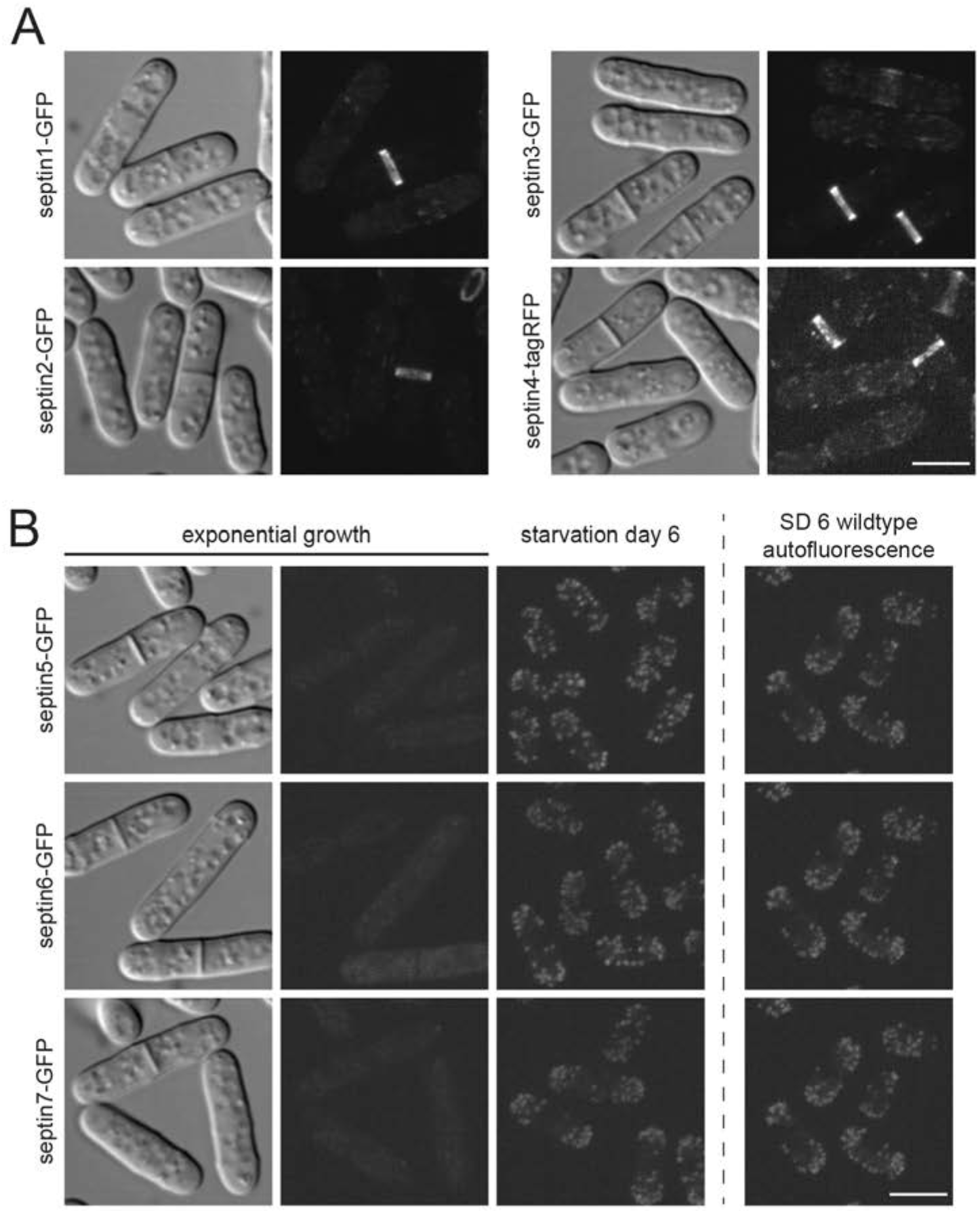
Localization of septins. (**A**) Spn1p-spn4p localization in exponential cells. (**B**) GFP-tagged spn5p, spn6p, and spn7p in exponential growth (left; corresponding DIC images show cell location) and on SD6 (middle). The unspecific signal portion can be estimated from comparison to the autofluorescence from a SD6 wild type cell without fluorescent tag with the same imaging and contrast settings (to the right of dashed line). Fluorescence images represent maximum intensity projections. Scale bars: 5µm.

**Figure 6 – figure supplement 1.**
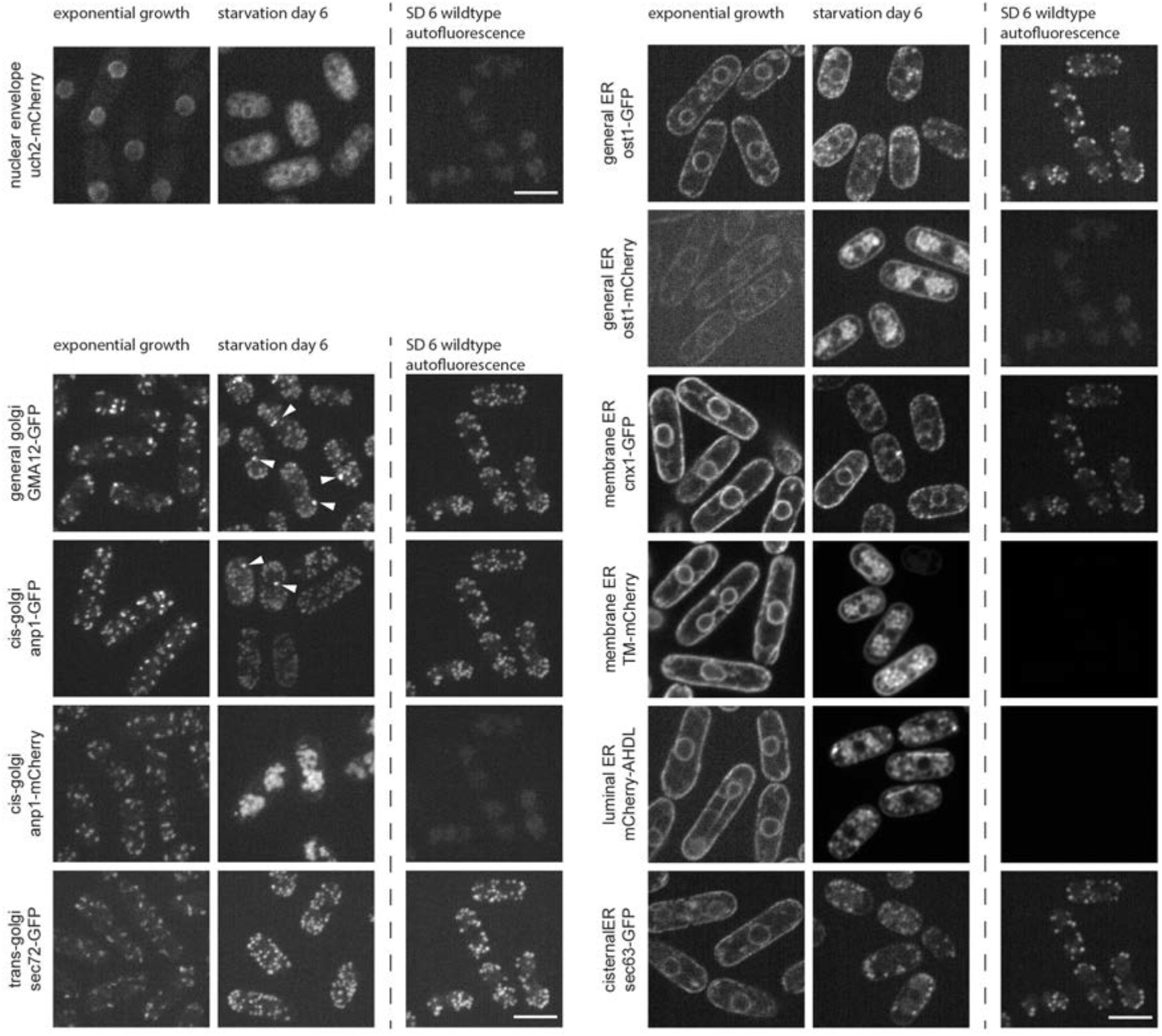
Subcellular architecture in deep starvation. (**A**) Images show fluorescence of markers for the indicated subcellular structures in exponentially growing cells and on SD6. The unspecific signal portion can be estimated from comparison to the autofluorescence from a SD6 wild type cell without fluorescent tag with the same imaging and contrast settings (to the right of dashed line). All images are maximum intensity projections. Scale bar: 5µm.

## Acknowledgements

We would like to thank Darren Gilmour, Stephen Huisman and Angela Lloyd for their critical reading of the manuscript. We would like to thank Craig Wang from Prof. Reinhard Furrer’s biostatistics group at the UZH for statistical consulting. We thank Mohan Balasubramanian, Kathleen L. Gould, Ian M. Hagan, Olaf Nielsen, Paul Nurse, Snezhana Oliferenko, John Pringle, Iva M. Tolic-Nørrelykke, Phong T. Tran, and Phillip D. Zamore for sharing strains. M.H. and C.I.N were supported by a program grant RGP0007–2010 from the Human Frontier Science Program to D.B, E.L.F., and A.H. M.B. was supported by a research grant from the Swiss National Science Foundation to DB. D.B. acknowledges funding from the University of Zurich.

## Movie legends

**Movie 1.** Motion arrest of lipid droplets in deep starvation

25sec DIC movies (4frames/sec) of cells in exponential growth and starvation days 2–8. Upper panels from left to right show cells in exponential growth and on starvation days 2, 3 and 4. Lowers panels show cells at starvation days 5, 6, 7 and 8. Scale bar: 5µm.

**Movie 2.** CF reversion after glucose addition

25sec DIC movies (4frames/sec) of cells on starvation day 5 before and after glucose addition. Upper panels from left to right show cells before glucose addition and 5min or 10min after glucose addition. Lower panels show cells 15min, 30min and 60min after glucose addition. Scale bar: 5µm.

**Movie 3.** Lipid droplet motion in cells under acute energy depletion

Left to right: 25sec DIC movies (4frames/sec) of cells exposed to acute glucose depletion and drug-induced energy depletion with 2hour and 0.5hour drug treatment respectively. Scale bar: 5µm.

**Movie 4.** Protoplasting of exponentially growing cells in hypertonic conditions

DIC movies (1frame/10sec) of protoplast evasion from the cell wall of exponentially growing cells in a dish. Cells are being incubated in 1.2M sorbitol containing cell wall digesting mix. Scale bar: 5µm.

**Movie 5.** Protoplasting of cells in hypotonic conditions

DIC movies (1frame/10sec) of cells, incubated on a dish with cell wall digesting enzymes in 0.5 M sorbitol containing buffer. Upper panels show from left to right, cells in exponential growth and on starvation day 6. The lowers panels show cells treated for acute glucose depletion and drug-induced energy depletion with 2hour and 0.5hour drug treatment respectively. Scale bar: 5µm.

**Movie 6.** F-actin organization in starvation

Movies (1frame/5sec) visualizing F-actin dynamics based on maximum intensity projections of Lifeact-GFP expressing cells. Upper panels from left to right show exponentially growing cells and cells at starvation days 2, 3 and 4. The lowers panels show cells at starvation days 5, 6, 7 and 8. Scale bar: 5µm.

**Movie 7.** CF in F-actin depleted cells

25sec DIC movies (4frames/sec) of wild type cells at starvation day 6 incubated from starvation day 3 onwards with DMSO (control, left panel) or LatB. Scale bar: 5µm.

**Movie 8.** Mitochondria dynamics during starvation

Movies (2frames/3sec) of mitochondria visualized by maximum intensity projections of cox4-GFP expressing cells during starvation. Panels show from left to right cells in exponential growth and cells on starvation days 3 and 6. Scale bar: 5µm.

**Movie 9.** Interference with autophagy delays CF

25sec DIC movies (4frames/sec) of from left to right, wild type, *atg1Δ* and *atg8Δ* cells at starvation days 6 (upper panel) and 9 (lower panel). Scale bar: 5µm.

## References

Abe, H., Shimoda, C., 2000. Autoregulated expression of Schizosaccharomyces pombe meiosis-specific transcription factor Mei4 and a genome-wide search for its target genes. Genetics 154, 1497–1508.

Adler, J., Parmryd, I., 2010. Quantifying colocalization by correlation: the Pearson correlation coefficient is superior to the Mander’s overlap coefficient. Cytometry A 77, 733–742. doi:10.1002/cyto.a.20896

Alvarez-Tabarés, I., Grallert, A., Ortiz, J.-M., Hagan, I.M., 2007. Schizosaccharomyces pombe protein phosphatase 1 in mitosis, endocytosis and a partnership with Wsh3/Tea4 to control polarised growth. Journal of Cell Science 120, 3589–3601. doi:10.1242/jcs.007567

An, H., Morrell, J.L., Jennings, J.L., Link, A.J., Gould, K.L., 2004. Requirements of fission yeast septins for complex formation, localization, and function. Mol. Biol. Cell 15, 5551–5564. doi:10.1091/mbc.E04-07-0640

Atilgan, E., Magidson, V., Khodjakov, A., Chang, F., 2015. Morphogenesis of the Fission Yeast Cell through Cell Wall Expansion. Curr. Biol. 25, 2150–2157. doi:10.1016/j.cub.2015.06.059

Bancaud, A., Huet, S., Rabut, G., Ellenberg, J., 2010. Fluorescence perturbation techniques to study mobility and molecular dynamics of proteins in live cells: FRAP, photoactivation, photoconversion, and FLIP. Cold Spring Harb Protoc 2010, 1303–1324. doi:10.1101/pdb.top90

Bähler, J., Wu, J.Q., Longtine, M.S., Shah, N.G., McKenzie, A., Steever, A.B., Wach, A., Philippsen, P., Pringle, J.R., 1998. Heterologous modules for efficient and versatile PCR-based gene targeting in Schizosaccharomyces pombe. Yeast 14, 943–951. doi:10.1002/(SICI)1097-0061(199807)14:10<943::AID-YEA292>3.0.CO;2-Y

Berlin, A., Paoletti, A., Chang, F., 2003. Mid2p stabilizes septin rings during cytokinesis in fission yeast. J. Cell Biol. 160, 1083–1092. doi:10.1083/jcb.200212016

Boke, E., Ruer, M., Wühr, M., Coughlin, M., Lemaitre, R., Gygi, S.P., Alberti, S., Drechsel, D., Hyman, A.A., Mitchison, T.J., 2016. Amyloid-like Self-Assembly of a Cellular Compartment. Cell 166, 637–650. doi:10.1016/j.cell.2016.06.051

Boothby, T.C., Tapia, H., Brozena, A.H., Piszkiewicz, S., Smith, A.E., Giovannini, I., Rebecchi, L., Pielak, G.J., Koshland, D., Goldstein, B., 2017. Tardigrades Use Intrinsically Disordered Proteins to Survive Desiccation. Molecular cell 65, 975–984.e5. doi:10.1016/j.molcel.2017.02.018

Brangwynne, C.P., Eckmann, C.R., Courson, D.S., Rybarska, A., Hoege, C., Gharakhani, J., Jülicher, F., Hyman, A.A., 2009. Germline P granules are liquid droplets that localize by controlled dissolution/condensation. Science 324, 1729–1732. doi:10.1126/science.1172046

Coelho, M., Dereli, A., Haese, A., Kühn, S., Malinovska, L., DeSantis, M.E., Shorter, J., Alberti, S., Gross, T., Tolić-Nørrelykke, I.M., 2013. Fission yeast does not age under favorable conditions, but does so after stress. Curr. Biol. 23, 1844–1852. doi:10.1016/j.cub.2013.07.084

Coelho, M., Lade, S.J., Alberti, S., Gross, T., Tolić, I.M., 2014. Fusion of protein aggregates facilitates asymmetric damage segregation. PLoS Biol. 12, e1001886. doi:10.1371/journal.pbio.1001886

Coller, H.A., Sang, L., Roberts, J.M., 2006. A new description of cellular quiescence. PLoS Biol. 4, e83. doi:10.1371/journal.pbio.0040083

Costantini, L.M., Baloban, M., Markwardt, M.L., Rizzo, M., Guo, F., Verkhusha, V.V., Snapp, E.L., 2015. A palette of fluorescent proteins optimized for diverse cellular environments. Nat Commun 6, 229–31. doi:10.1038/ncomms8670

Costello, G., Rodgers, L., Beach, D., 1986. Fission yeast enters the stationary phase G0 state from either mitotic G1 or G2. Current Genetics 2, 119–125. doi:10.1007/BF00378203

Cowan, A.E., Koppel, D.E., Setlow, B., Setlow, P., 2003. A soluble protein is immobile in dormant spores of Bacillus subtilis but is mobile in germinated spores: implications for spore dormancy. Proc Natl Acad Sci USA 100, 4209–4214. doi:10.1073/pnas.0636762100

Dijksterhuis, J., Nijsse, J., Hoekstra, F.A., Golovina, E.A., 2007. High viscosity and anisotropy characterize the cytoplasm of fungal dormant stress-resistant spores. Eukaryotic Cell 6, 157–170. doi:10.1128/EC.00247-06

Dunn, K.W., Kamocka, M.M., McDonald, J.H., 2011. A practical guide to evaluating colocalization in biological microscopy. Am. J. Physiol., Cell Physiol. 300, C723–42. doi:10.1152/ajpcell.00462.2010

Egel, R., 1989. Mating-Type Genes, Meiosis, and Sporulation, in: Molecular Biology of the Fission Yeast. Elsevier, pp. 31–73. doi:10.1016/B978-0-12-514085-0.50007-5

Elbein, A.D., Pan, Y.T., Pastuszak, I., Carroll, D., 2003. New insights on trehalose: a multifunctional molecule. Glycobiology 13, 17R–27R. doi:10.1093/glycob/cwg047

Fels, J., Orlov, S.N., Grygorczyk, R., 2009. The hydrogel nature of mammalian cytoplasm contributes to osmosensing and extracellular pH sensing. Biophys. J. 96, 4276–4285. doi:10.1016/j.bpj.2009.02.038

Fu, C., Jain, D., Costa, J., Velve-Casquillas, G., Tran, P.T., 2011. mmb1p binds mitochondria to dynamic microtubules. Curr. Biol. 21, 1431–1439. doi:10.1016/j.cub.2011.07.013

Fullerton, G.D., Kanal, K.M., Cameron, I.L., 2006. Osmotically unresponsive water fraction on proteins: non-ideal osmotic pressure of bovine serum albumin as a function of pH and salt concentration. Cell Biol. Int. 30, 86–92. doi:10.1016/j.cellbi.2005.11.001

Grygorczyk, R., Boudreault, F., Platonova, A., Orlov, S.N., 2015. Salt and osmosensing: role of cytoplasmic hydrogel. Pflugers Arch. 467, 475–487. doi:10.1007/s00424-014-1680-2

Guppy, M., Withers, P., 1999. Metabolic depression in animals: physiological perspectives and biochemical generalizations. Biol Rev Camb Philos Soc 74, 1–40.

Han, T.W., Kato, M., Xie, S., Wu, L.C., Mirzaei, H., Pei, J., Chen, M., Xie, Y., Allen, J., Xiao, G., McKnight, S.L., 2012. Cell-free formation of RNA granules: bound RNAs identify features and components of cellular assemblies. Cell 149, 768–779. doi:10.1016/j.cell.2012.04.016

Huang, J., Huang, Y., Yu, H., Subramanian, D., Padmanabhan, A., Thadani, R., Tao, Y., Tang, X., Wedlich-Soldner, R., Balasubramanian, M.K., 2012. Nonmedially assembled F-actin cables incorporate into the actomyosin ring in fission yeast. J. Cell Biol. 199, 831–847. doi:10.1083/jcb.201209044

Huh, W.-K., Falvo, J.V., Gerke, L.C., Carroll, A.S., Howson, R.W., Weissman, J.S., O’Shea, E.K., 2003. Global analysis of protein localization in budding yeast. Nature 425, 686–691. doi:10.1038/nature02026

Jain, A., Vale, R.D., 2017. RNA phase transitions in repeat expansion disorders. Nature 546, 243–247. doi:10.1038/nature22386

Jankovics, F., Brunner, D., 2006. Transiently Reorganized Microtubules Are Essential for Zippering during Dorsal Closure in Drosophila melanogaster. Dev. Cell 11, 375–385. doi:10.1016/j.devcel.2006.07.014

Joyner, R.P., Tang, J.H., Helenius, J., Dultz, E., Brune, C., Holt, L.J., Huet, S., Müller, D.J., Weis, K., 2016. A glucose-starvation response regulates the diffusion of macromolecules. Elife 5, e09376. doi:10.7554/eLife.09376

Kato, M., Han, T.W., Xie, S., Shi, K., Du, X., Wu, L.C., Mirzaei, H., Goldsmith, E.J., Longgood, J., Pei, J., Grishin, N.V., Frantz, D.E., Schneider, J.W., Chen, S., Li, L., Sawaya, M.R., Eisenberg, D., Tycko, R., McKnight, S.L., 2012. Cell-free formation of RNA granules: low complexity sequence domains form dynamic fibers within hydrogels. Cell 149, 753–767. doi:10.1016/j.cell.2012.04.017

Kelly, F.D., Nurse, P., 2011. De novo growth zone formation from fission yeast spheroplasts. PLoS ONE 6, e27977. doi:10.1371/journal.pone.0027977

Kim, D.U., Hayles, J., Kim, D., Wood, V., Park, H.-O., Won, M., Yoo, H.-S., Duhig, T., Nam, M., Palmer, G., Han, S., Jeffery, L., Baek, S.-T., Lee, H., Shim, Y.S., Lee, M., Kim, L., Heo, K.-S., Noh, E.J., Lee, A.-R., Jang, Y.-J., Chung, K.-S., Choi, S.-J., Park, J.-Y., Park, Y., Kim, H.M., Park, S.-K., Park, H.-J., Kang, E.-J., Kim, H.B., Kang, H.-S., Park, H.-M., Kim, K., Song, K., Song, K.B., Nurse, P., Hoe, K.-L., 2010. Analysis of a genome-wide set of gene deletions in the fission yeast Schizosaccharomyces pombe. Nat. Biotechnol. 28, 617–623. doi:10.1038/nbt.1628

Laporte, D., Courtout, F., Pinson, B., Dompierre, J., Salin, B., Brocard, L., Sagot, I., 2015. A stable microtubule array drives fission yeast polarity reestablishment upon quiescence exit. J. Cell Biol. 210, 99–113. doi:10.1083/jcb.201502025

Laporte, D., Salin, B., Daignan-Fornier, B., 2008. Reversible cytoplasmic localization of the proteasome in quiescent yeast cells. The Journal of cell 181, 737–745. doi:10.1083/jcb.200711154

Lennon, J.T., Jones, S.E., 2011. Microbial seed banks: the ecological and evolutionary implications of dormancy. Nat. Rev. Microbiol. 9, 119–130. doi:10.1038/nrmicro2504

Leupold, U., 1950. Die Vererbung von Homothallie und Heterothallie bei Schizosaccharomyces pombe, C. R. Trav. Lab. Carlsberg Ser. Physiol.

Listenberger, L.L., Brown, D.A., 2007. Fluorescent detection of lipid droplets and associated proteins. Curr Protoc Cell Biol Chapter 24, Unit 24.2–24.2.11. doi:10.1002/0471143030.cb2402s35

Long, A.P., Manneschmidt, A.K., VerBrugge, B., Dortch, M.R., Minkin, S.C., Prater, K.E., Biggerstaff, J.P., Dunlap, J.R., Dalhaimer, P., 2012. Lipid droplet de novo formation and fission are linked to the cell cycle in fission yeast. Traffic 13, 705–714. doi:10.1111/j.1600-0854.2012.01339.x

Longtine, M.S., DeMarini, D.J., Valencik, M.L., Al-Awar, O.S., Fares, H., De Virgilio, C., Pringle, J.R., 1996. The septins: roles in cytokinesis and other processes. Curr. Opin. Cell Biol. 8, 106–119. doi:10.1016/S0955-0674(96)80054-8

Luby-Phelps, K., Taylor, D.L., Lanni, F., 1986. Probing the structure of cytoplasm. J. Cell Biol. 102, 2015–2022.

Makushok, T., Alves, P., Huisman, S.M., Kijowski, A.R., Brunner, D., 2016. Sterol-Rich Membrane Domains Define Fission Yeast Cell Polarity. Cell 165, 1182–1196. doi:10.1016/j.cell.2016.04.037

Mata, J., Lyne, R., Burns, G., Bähler, J., 2002. The transcriptional program of meiosis and sporulation in fission yeast. Nat. Genet. 32, 143–147. doi:10.1038/ng951

McDonald, K., Schwarz, H., Müller-Reichert, T., Webb, R., Buser, C., Morphew, M., 2010. “Tips and Tricks” for High-Pressure Freezing of Model Systems, in: Electron Microscopy of Model Systems, Methods in Cell Biology. Elsevier, pp. 671–693. doi:10.1016/S0091-679X(10)96028-7

Miermont, A., Waharte, F., Hu, S., McClean, M.N., Bottani, S., Leon, S., Hersen, P., 2013. Severe osmotic compression triggers a slowdown of intracellular signaling, which can be explained by molecular crowding. Proc Natl Acad Sci USA 110, 5725–5730. doi:10.1073/pnas.1215367110

Mitchison, T.J., Charras, G.T., Mahadevan, L., 2008. Implications of a poroelastic cytoplasm for the dynamics of animal cell shape. Seminars in Cell & Developmental Biology 19, 215–223. doi:10.1016/j.semcdb.2008.01.008

Moeendarbary, E., Valon, L., Fritzsche, M., Harris, A.R., Moulding, D.A., Thrasher, A.J., Stride, E., Mahadevan, L., Charras, G.T., 2013. The cytoplasm of living cells behaves as a poroelastic material. Nat Mater 12, 253–261. doi:10.1038/nmat3517

Moreno, S., Klar, A., Nurse, P., 1991. Molecular genetic analysis of fission yeast Schizosaccharomyces pombe. Meth. Enzymol. 194, 795–823. doi:10.1016/0076-6879(91)94059-l

Mostowy, S., Cossart, P., 2012. Septins: the fourth component of the cytoskeleton. Nat. Rev. Mol. Cell Biol. 13, 183–194. doi:10.1038/nrm3284

Munder, M.C., Midtvedt, D., Franzmann, T., Nüske, E., Otto, O., Herbig, M., Ulbricht, E., Müller, P., Taubenberger, A., Maharana, S., Malinovska, L., Richter, D., Guck, J., Zaburdaev, V., Alberti, S., 2016. A pH-driven transition of the cytoplasm from a fluid-to a solid-like state promotes entry into dormancy. Elife 5, e09347. doi:10.7554/eLife.09347

Nakatogawa, H., Suzuki, K., Kamada, Y., Ohsumi, Y., 2009. Dynamics and diversity in autophagy mechanisms: lessons from yeast. Nat. Rev. Mol. Cell Biol. 10, 458–467. doi:10.1038/nrm2708

Narayanaswamy, R., Levy, M., Tsechansky, M., Stovall, G.M., O’Connell, J.D., Mirrielees, J., Ellington, A.D., Marcotte, E.M., 2009. Widespread reorganization of metabolic enzymes into reversible assemblies upon nutrient starvation. Proc. Natl. Acad. Sci. U.S.A. 106, 10147–10152. doi:10.1073/pnas.0812771106

Noda, T., 2008. Chapter 2 Viability Assays to Monitor Yeast Autophagy, in: Autophagy: Lower Eukaryotes and Non-Mammalian Systems, Part A, Methods in Enzymology. Elsevier, pp. 27–32. doi:10.1016/S0076-6879(08)03202-3

Noree, C., Sato, B.K., Broyer, R.M., Wilhelm, J.E., 2010. Identification of novel filament-forming proteins in Saccharomyces cerevisiae and Drosophila melanogaster. J. Cell Biol. 190, 541–551. doi:10.1083/jcb.201003001

O’Connell, J.D., Zhao, A., Ellington, A.D., Marcotte, E.M., 2012. Dynamic reorganization of metabolic enzymes into intracellular bodies. Annu. Rev. Cell Dev. Biol. 28, 89–111. doi:10.1146/annurevcellbio-101011-155841

Oda, A., Takemata, N., Hirata, Y., Miyoshi, T., Suzuki, Y., Sugano, S., Ohta, K., 2015. Dynamic transition of transcription and chromatin landscape during fission yeast adaptation to glucose starvation. Genes Cells 20, 392–407. doi:10.1111/gtc.12229

Onishi, M., Koga, T., Hirata, A., Nakamura, T., Asakawa, H., Shimoda, C., Bähler, J., Wu, J.-Q., Takegawa, K., Tachikawa, H., Pringle, J.R., Fukui, Y., 2010. Role of septins in the orientation of forespore membrane extension during sporulation in fission yeast. Mol. Cell. Biol. 30, 2057–2074. doi:10.1128/MCB.01529-09

Pan, F., Malmberg, R.L., Momany, M., 2007. Analysis of septins across kingdoms reveals orthology and new motifs. BMC Evol. Biol. 7, 103. doi:10.1186/1471-2148-7-103

Pardo, M., Nurse, P., 2005. The nuclear rim protein Amo1 is required for proper microtubule cytoskeleton organisation in fission yeast. Journal of Cell Science 118, 1705–1714. doi:10.1242/jcs.02305

Parry, B.R., Surovtsev, I.V., Cabeen, M.T., O’Hern, C.S., Dufresne, E.R., Jacobs-Wagner, C., 2014. The bacterial cytoplasm has glass-like properties and is fluidized by metabolic activity. Cell 156, 183–194. doi:10.1016/j.cell.2013.11.028

Patel, A., Lee, H.O., Jawerth, L., Maharana, S., Jahnel, M., Hein, M.Y., Stoynov, S., Mahamid, J., Saha, S., Franzmann, T.M., Pozniakovski, A., Poser, I., Maghelli, N., Royer, L.A., Weigert, M., Myers, E.W., Grill, S., Drechsel, D., Hyman, A.A., Alberti, S., 2015. A Liquid-to-Solid Phase Transition of the ALS Protein FUS Accelerated by Disease Mutation. Cell 162, 1066–1077. doi:10.1016/j.cell.2015.07.047

Petrovska, I., Nüske, E., Munder, M.C., Kulasegaran, G., Malinovska, L., Kroschwald, S., Richter, D., Fahmy, K., Gibson, K., Verbavatz, J.-M., Alberti, S., 2014. Filament formation by metabolic enzymes is a specific adaptation to an advanced state of cellular starvation. Elife 3, e02409. doi:10.7554/eLife.02409

Reuner, A., Hengherr, S., Mali, B., Förster, F., Arndt, D., Reinhardt, R., Dandekar, T., Frohme, M., Brümmer, F., Schill, R.O., 2009. Stress response in tardigrades: differential gene expression of molecular chaperones. Cell Stress and Chaperones 15, 423–430. doi:10.1007/s12192-009-0158-1

Riedl, J., Crevenna, A.H., Kessenbrock, K., Yu, J.H., Neukirchen, D., Bista, M., Bradke, F., Jenne, D., Holak, T.A., Werb, Z., Sixt, M., Wedlich-Soldner, R., 2008. Lifeact: a versatile marker to visualize F-actin. Nature methods 5, 605–607. doi:10.1038/nmeth.1220

Sagot, I., Pinson, B., Salin, B., Daignan-Fornier, B., 2006. Actin bodies in yeast quiescent cells: an immediately available actin reserve? Mol. Biol. Cell 17, 4645–4655. doi:10.1091/mbc.E06-04-0282

Saitoh, S., Yanagida, M., 2014. Does a shift to limited glucose activate checkpoint control in fission yeast? FEBS Letters 588, 2373–2378. doi:10.1016/j.febslet.2014.04.047

Sbalzarini, I.F., Koumoutsakos, P., 2005. Feature point tracking and trajectory analysis for video imaging in cell biology. J. Struct. Biol. 151, 182–195. doi:10.1016/j.jsb.2005.06.002

Sigova, A., Rhind, N., Zamore, P.D., 2004. A single Argonaute protein mediates both transcriptional and posttranscriptional silencing in Schizosaccharomyces pombe. Genes Dev. 18, 2359–2367. doi:10.1101/gad.1218004

Snapp, E.L., 2003. Formation of stacked ER cisternae by low affinity protein interactions. J. Cell Biol. 163, 257–269. doi:10.1083/jcb.200306020

Soto, T., Fernandez, J., Vicente-Soler, J., Cansado, J., Gacto, M., 1999. Accumulation of trehalose by overexpression of tps1, coding for trehalose-6-phosphate synthase, causes increased resistance to multiple stresses in the fission yeast schizosaccharomyces pombe. Appl. Environ. Microbiol. 65, 2020–2024.

Spector, I., Shochet, N.R., Kashman, Y., Groweiss, A., 1983. Latrunculins: novel marine toxins that disrupt microfilament organization in cultured cells. Science 219, 493–495.

Storey, K.B., Storey, J.M., 2007. Tribute to P. L. Lutz: putting life on “pause” – molecular regulation of hypometabolism. J. Exp. Biol. 210, 1700–1714. doi:10.1242/jeb.02716

Sun, W.Q., Leopold, A.C., 1997. Cytoplasmic Vitrification and Survival of Anhydrobiotic Organisms. Comparative Biochemistry and Physiology Part A: Physiology 117, 327–333. doi:10.1016/S0300-9629(96)00271-X

Tamai, Y., Tanaka, H., Nakanishi, K., 1996. Molecular Dynamics Study of Polymer−Water Interaction in Hydrogels. 1. Hydrogen-Bond Structure. Macromolecules 29, 6750–6760. doi:10.1021/ma951635z

Tanaka, K., Hirata, A., 1982. Ascospore development in the fission yeasts Schizosaccharomyces pombe and S. japonicus. Journal of Cell Science 56, 263–279.

Tasto, J.J., Morrell, J.L., Gould, K.L., 2003. An anillin homologue, Mid2p, acts during fission yeast cytokinesis to organize the septin ring and promote cell separation. J. Cell Biol. 160, 1093–1103. doi:10.1083/jcb.200211126

Tolić-Nørrelykke, I.M., Munteanu, E.-L., Thon, G., Oddershede, L., Berg-Sørensen, K., 2004. Anomalous diffusion in living yeast cells. Phys. Rev. Lett. 93, 078102. doi:10.1103/PhysRevLett.93.078102

Vejrup-Hansen, R., Fleck, O., Landvad, K., Fahnøe, U., Broendum, S.S., Schreurs, A.-S., Kragelund, B.B., Carr, A.M., Holmberg, C., Nielsen, O., 2014. Spd2 assists Spd1 in the modulation of ribonucleotide reductase architecture but does not regulate deoxynucleotide pools. Journal of Cell Science 127, 2460–2470. doi:10.1242/jcs.139816

Vjestica, A., Tang, X.-Z., Oliferenko, S., 2008. The actomyosin ring recruits early secretory compartments to the division site in fission yeast. Mol. Biol. Cell 19, 1125–1138. doi:10.1091/mbc.E07-07-0663

Wang, H., Tang, X., Liu, J., Trautmann, S., Balasundaram, D., McCollum, D., Balasubramanian, M.K., 2002. The multiprotein exocyst complex is essential for cell separation in Schizosaccharomyces pombe. Mol. Biol. Cell 13, 515–529. doi:10.1091/mbc.01-11-0542

Watanabe, T., Miyashita, K., Saito, T.T., Yoneki, T., Kakihara, Y., Nabeshima, K., Kishi, Y.A., Shimoda, C., Nojima, H., 2001. Comprehensive isolation of meiosis-specific genes identifies novel proteins and unusual non-coding transcripts in Schizosaccharomyces pombe. Nucleic Acids Res. 29, 2327–2337.

Weiss, M., Elsner, M., Kartberg, F., Nilsson, T., 2004. Anomalous subdiffusion is a measure for cytoplasmic crowding in living cells. Biophys. J. 87, 3518–3524. doi:10.1529/biophysj.104.044263

Wirtz, D., 2009. Particle-tracking microrheology of living cells: principles and applications. Annu Rev Biophys 38, 301–326. doi:10.1146/annurev.biophys.050708.133724

Wu, J.-Q., Ye, Y., Wang, N., Pollard, T.D., Pringle, J.R., 2010. Cooperation between the septins and the actomyosin ring and role of a cell-integrity pathway during cell division in fission yeast. Genetics 186, 897–915. doi:10.1534/genetics.110.119842

Yaffe, M.P., Stuurman, N., Vale, R.D., 2003. Mitochondrial positioning in fission yeast is driven by association with dynamic microtubules and mitotic spindle poles. Proc Natl Acad Sci USA 100, 11424–11428. doi:10.1073/pnas.1534703100

Yanagida, M., 2009. Cellular quiescence: are controlling genes conserved? Trends in Cell Biology 19, 705–715. doi:10.1016/j.tcb.2009.09.006

Yanagida, M., Ikai, N., Shimanuki, M., Sajiki, K., 2011. Nutrient limitations alter cell division control and chromosome segregation through growth-related kinases and phosphatases. Philos. Trans. R. Soc. Lond., B, Biol. Sci. 366, 3508–3520. doi:10.1098/rstb.2011.0124

Yang, F., Moss, L.G., Phillips, G.N., 1996. The molecular structure of green fluorescent protein. Nat. Biotechnol. 14, 1246–1251. doi:10.1038/nbt1096-1246

Zhang, D., Vjestica, A., Oliferenko, S., 2012. Plasma membrane tethering of the cortical ER necessitates its finely reticulated architecture. Curr. Biol. 22, 2048–2052. doi:10.1016/j.cub.2012.08.047

Zhang, D., Vjestica, A., Oliferenko, S., 2010. The Cortical ER Network Limits the Permissive Zone for Actomyosin Ring Assembly. Current Biology 20, 1029–1034. doi:10.1016/j.cub.2010.04.017

